# Identification of potential SARS-CoV-2 genetic markers resulting from host domestication

**DOI:** 10.1101/2024.07.27.605454

**Authors:** Janusz Wiśniewski, Heng-Chang Chen

## Abstract

We developed a *k*-mer-based pipeline, namely the Pathogen Origin Recognition Tool using Enriched *K*-mers (PORT-EK) to identify genomic regions enriched in the respective hosts after the comparison of metagenomes of isolates between two host species. Using it we identified thousands of *k*-mers enriched in US white-tailed deer and betacoronaviruses in bat reservoirs while comparing them with human isolates. We demonstrated different coverage landscapes of *k*-mers enriched in deer and bats and unraveled 148 mutations in enriched *k*-mers yielded from the comparison of viral metagenomes between bat and human isolates. We observed that the third position within a genetic codon is prone to mutations, resulting in a high frequency of synonymous mutations of amino acids harboring the same physicochemical properties as unaltered amino acids. Finally, we classified and predicted the likelihood of host species based on the enriched *k*-mer counts. Altogether, PORT-EK showcased its feasibility for identifying enriched viral genomic regions, illuminating the different intrinsic tropisms of coronavirus after host domestication.

**Teaser:** A measure of enriched viral genomic correlates resulting from host domestication as a potential predictor of zoonotic risk.

## Introduction

It has been estimated that nearly two-thirds of emerging infectious diseases that affect humans have their origins in animal reservoirs, namely zoonoses (*1*). To date, at least 20 zoonoses have been discovered in Europe, and 14 have spread worldwide, posing potential risks that may threaten public health. Coronaviruses are a classic example of zoonotic viruses that infect humans. The first outbreak of a coronavirus-caused epidemic (severe acute respiratory syndrome caused by SARS-CoV) in humans can be tracked to 2002-2003 (*2-4*). The recent COVID-19 pandemic (*5*, *6*) alerted us to the importance of rapidly controlling the spread of infectious zoonotic diseases and their prevention.

Like other RNA viruses, coronaviruses are genetically heterogeneous, in part due to the highly error-prone and low-fidelity RNA-dependent RNA polymerase (RdRP) that replicates their genomes (*7-9*). In addition to RdRP, SARS-CoV-2 possesses an associated proofreading enzyme, exoribonuclease. The discontinuous nature of coronavirus transcription results in coronaviruses with high rates of recombination, insertions, deletions, and point mutations (*10*), thereby enabling frequent host switching under different selection pressures. As such, understanding the potential viral genomic correlates underpinning host jumps in viruses across humans and other animal reservoirs is essential to mitigating viral threats across species boundaries.

Humans are just one of many intermediate or recipient hosts within a large and complex network of host reservoirs where the unhindered transmission of zoonotic viruses can be achieved. However, relative to studies focusing on the mechanistic interplay between SARS-CoV-2 and humans, viral cross-species transmission between animal hosts and intrahost viral circulation within animal reservoirs remains understudied. Extending these investigations toward the broader evolutionary processes aligning with host jumps across animal hosts may also enhance our understanding of SARS-CoV-2 pathogenesis and reinforce both human and animal health in the fight against zoonotic infections.

Indeed, after the outbreak of the COVID-19 pandemic, the vast majority of studies paid attention to viral genetic diversity and the consequence of viral transmission and pathogenesis; however, our current understanding remains insufficient to effectively predict, prevent, and manage current and imminent zoonotic disease threats. One of the reasons for this is the biased focus on investigating the structural proteins and associated mutations of SARS-CoV-2 at the protein level instead of an overall measurement of the whole viral genome. In addition, most studies do not include genomic data in their analyses despite the fact that genomic analyses are critical for investigating the drivers of viral host jumps (*11*). For this reason, in this study, we examine the entirety of available metagenomic sequences of isolates collected from different SARS-CoV-2 host species to identify the genomic regions over-represented in respective host species.

Methodologies using *k-*mers representation directly compare the counts of nucleotide sequences of the length *k* between samples (*12*, *13*). A common step is to break a reference sequence into *k*-mers and use them to create a hash table. In parallel, target sequences are broken into *k*-mers and queried against the hash table to check for shared *k*-mers (*14-17*). Different *k*-mer-based models have been developed to optimize sequence analysis and comparison, such as the *k*-mer sparse matrix model for sequence comparison (*18*) and an anomaly detection algorithm (*19*, *20*). These approaches have gained momentum in high-throughput sequencing data analysis (*21-25*) and have been central to the field of metagenomics, where they are used to discover unique *k*-mer signatures to classify organisms (*26, 27*) and capture biological variations and functional annotation in RNA-seq data (*12*, *28*). More recently, the *k*-mer counting strategy has also been applied to identify genomic sequence signatures harboring mutations across thousands of SARS-CoV-2 genomic sequences (*29*).

In this work, to deepen our grasp on the viral genomic diversity associated with host domestication, we present a *k*-mer-based approach, namely the Pathogen Origin Recognition Tool using Enriched *K*-mers (PORT-EK), which allows for a comparison of different metagenomic datasets encompassing viral genomic sequences captured from two host species and the identification of over-represented genomic regions, *k*- mers, correlated to specific hosts. We generated three metagenomic datasets: the deer, bat, and out of the bag (OoB) datasets. Each dataset consists of genomic sequences of SARS-CoV-2 isolates from two different hosts, allowing for a comparison of the enrichment of *k*-mers between two species. In the deer dataset, we utilized SARS-CoV-2 genomic sequences isolated from US white-tailed deer. The genomic sequences of deer isolates present in this dataset originate from one single host, US white-tailed deer, offering an ideal model to dissect intrahost circulation with a continuous time frame inside an enclosed geography. In addition for the identification of over-represented viral genomic regions in respective host species, we applied the enriched *k*-mer counts computed based on these three metagenomic datasets for the classification and prediction of the most probable host species of a viral genomic sequence, highlighting the potential that enriched *k-*mers could serve as genetic markers for predicting the likelihood of distinct host reservoirs of SARS-CoV-2.

## Results

### Design and analytical pipeline of PORT-EK

The principle of *k*-mer-based approaches is to count the number of distinct substrings of length *k* in a string *S*, or a set of strings, where *k* is a positive integer (*30*) representing a unique mark in an indicated string *S*. Once we determined the length of *k* (see the following paragraph), we broke the whole SARS-CoV-2 genomic sequences down to 15 nt sequences, namely 15 nt *k*- mers with overlaps, and compared the counts of 15 nt *k*-mers between samples. The workflow of PORT-EK consists of four main blocks, including (1) *k*-mers matrix preparation, (2) *k*-mers filtering and selection, (3) identification of mutations in enriched *k*-mers, and (4) classification of the likelihood of hosts based on the enriched *k*-mer counts (**Fig. 1A**). We detail each step in PORT-EK as follows: First, we extract all *k*-mers of length *k* from metagenomic datasets and record their counts. At this stage, we discard the *k*-mers with any ambiguous nucleotides (first layer of filtering) and construct a count matrix containing counts of all non-redundant *k*-mers from all input metagenomic sequences. Second, we calculate the frequency of appearance of *k*-mers (detailed in **Materials and Method**) and the average counts of all filtered *k*-mers present in the same host isolates. We keep *k*-mers that pass through a tunable frequency threshold and mark them as “common” *k*-mers; for those that fail, we mark them as “rare” *k*-mers (**Fig. 1B**; second layer of filtering). Third, we compute the difference in the average count of *k*-mers and root mean square error (RMSE, detailed in **Materials and Method**) between the host isolates (third layer of filtering). Of note, it is the step at which the optimal length of *k*-mers is determined. From a pool of the common *k*-mers, we retrieve those with statistical significance, showing differences in the average count between two host isolates. They are the *k*- mers over-represented in the isolates from the same host (**Fig. 1B**). In parallel, we inspect the sequence of rare *k*-mers and pool them with common *k*-mers when sequences of rare *k*-mers resemble any common *k*- mer. A mismatch of a maximum of two nucleotides between two sequences (rare versus common *k*-mers) is allowed. Finally, to obtain the final output of the *k*-mers, we once again apply an RMSE-based filter to those *k*-mers over-represented in isolates from the same host, enabling the concentration of enriched *k*-mers identified in one specific host (the fourth layer of filtering). The hieratical filtering procedure used for the identification of enriched *k*-mers is illustrated in **Figure 1B**.

**Fig. 1.**
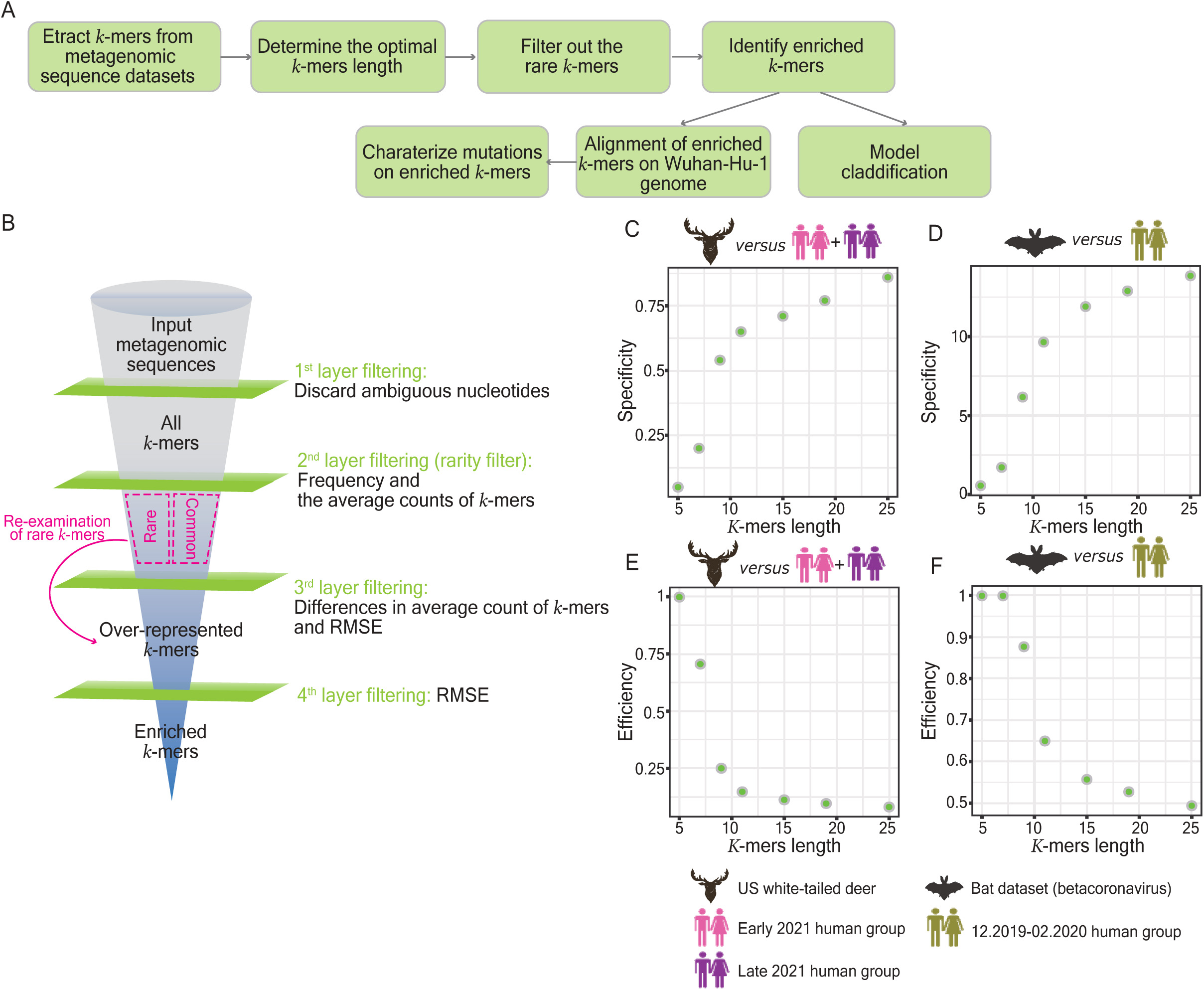
Rational design of PORT-EK and determination of the enriched *k*-mers. (**A**) The analytical pipeline of PORT-EK. PORT-EK consists of four steps including (1) *k*-mers matrix preparation, (2) *k*-mers filtering and selection, (3) the identification of host-specific mutations, and (4) the classification of hosts. Details are described in the main text. (**B**) Funnel plot representing the filtering strategies for the selection of enriched *k*-mers. Four layers of filtering were implemented in the PORT-EK pipeline. (**C**, **D**) Dot plots representing the specificity among *k*-mers with different lengths based on the calculation of the mean *k*- mers counts RMSE of common *k*-mers retrieved in the deer (**C**) and bat (**D**) datasets, respectively. (**E**, **F**) Dot plots representing the computational efficiency based on the percentage of the *k*-mers passing through the rarity filter in the deer (**E**) and bat (**F**) datasets to determine the optimal length of *k*-mers with affordable computing time and memory size. Details of the calculation of the specificity and the computational efficiency are described in **Materials and Methods**. Overall, the *k*-mers of 15 nt in length were considered to be optimal and chosen for this work.

### Determination of the optimal length of *k*-mers using SARS-CoV-2 isolates in the deer and bat datasets

Once the analytical pipeline of PORT-EK is optimized, our first task is to determine the optimal length of *k*- mers that can demonstrate the maximum specificity of enrichment between different host species using our collected metagenomics data. To compute enriched *k*-mers, we analyzed 336 SARS-CoV-2 genomic sequences in deer isolates compared to human isolates in April 2021 (n = 21,906, namely the early 2021 human group) and in November 2021 (n = 11,525, namely late 2021 human group), respectively. With respect to SARS-CoV-2 isolates in bats (n = 263), we analyzed their genomic sequences in contrast with those from humans between December 2019 and February 2020 (n = 2081). The datasets are listed in **Supplementary Tables S1-S3**, **S6** and **S7**. Given the geographical differences and inconsistent sample collection dates between isolates in both datasets, we separated SARS-CoV-2 isolates from humans from those geographically and temporally aligned with sequences isolated from both animal reservoirs.

We examined *k*-mers in the length of 5 nt, 7 nt, 11 nt, 15 nt, 19 nt, and 25 nt, followed by the calculation of the average count of *k*-mers and RMSE, representing the specificity of the *k*-mers. Here, a plateau of saturation was observed, starting at the *k-*mer length approaching 15 nt in both datasets (**Fig. 1C** and **1D**). It is important to note that although the RMSE increases as the length of the *k*-mers extends to 19 and 25 nt, we assume that longer *k*-mers may encounter a higher probability of being biased because of the genomic variations in different coronavirus variants, such as deletions and recombinations, thereby masking the intrinsic difference of the genomic properties due to *k*-mer decomposition into shorter pieces. In addition, while computing the efficiency of *k*-mer extraction, as represented by the ratio between common and rare *k-* mers of different lengths, we observed that *k*-mers of 15 nt in length offer optimal efficiency in the context of affordable computing power and memory size in both animal datasets (**Fig. 1E** and **1F**). Based on the calculation of *k*-mer specificity and efficiency, we chose *k*-mers of 15 nt in length for the further characterization of enriched *k*-mers.

### Identification of enriched *k*-mers in SARS-CoV-2 isolates in deer and bat datasets

It is important to stress that, using PORT-EK, we aim to seek any viral genomic region that is quantitatively enriched in one host. Thus, we calculated the difference in the average count of individual *k-*mers between two host species (**Fig. 1B**, the third layer of filtering) and tested their statistical significance designated as its *p*-value (detailed in **Materials and Methods**) in the deer (**Fig. 2A-2C**) and bat datasets (**Fig. 2D**). For *k*-mers over-represented in deer isolates, we only retrained the ones that were over-represented against both the early and late 2021 human groups to achieve a greater probability that the identified *k*-mers result from a consecutive period of intrahost selection pressure (**Fig. 2C**). We cannot perform this selection on the *k*-mers retrieved in the bat dataset because no clear temporal boundary exists in relation to bat isolates restricted in the same geographical regions nor in human isolates used in the bat dataset.

**Fig. 2.**
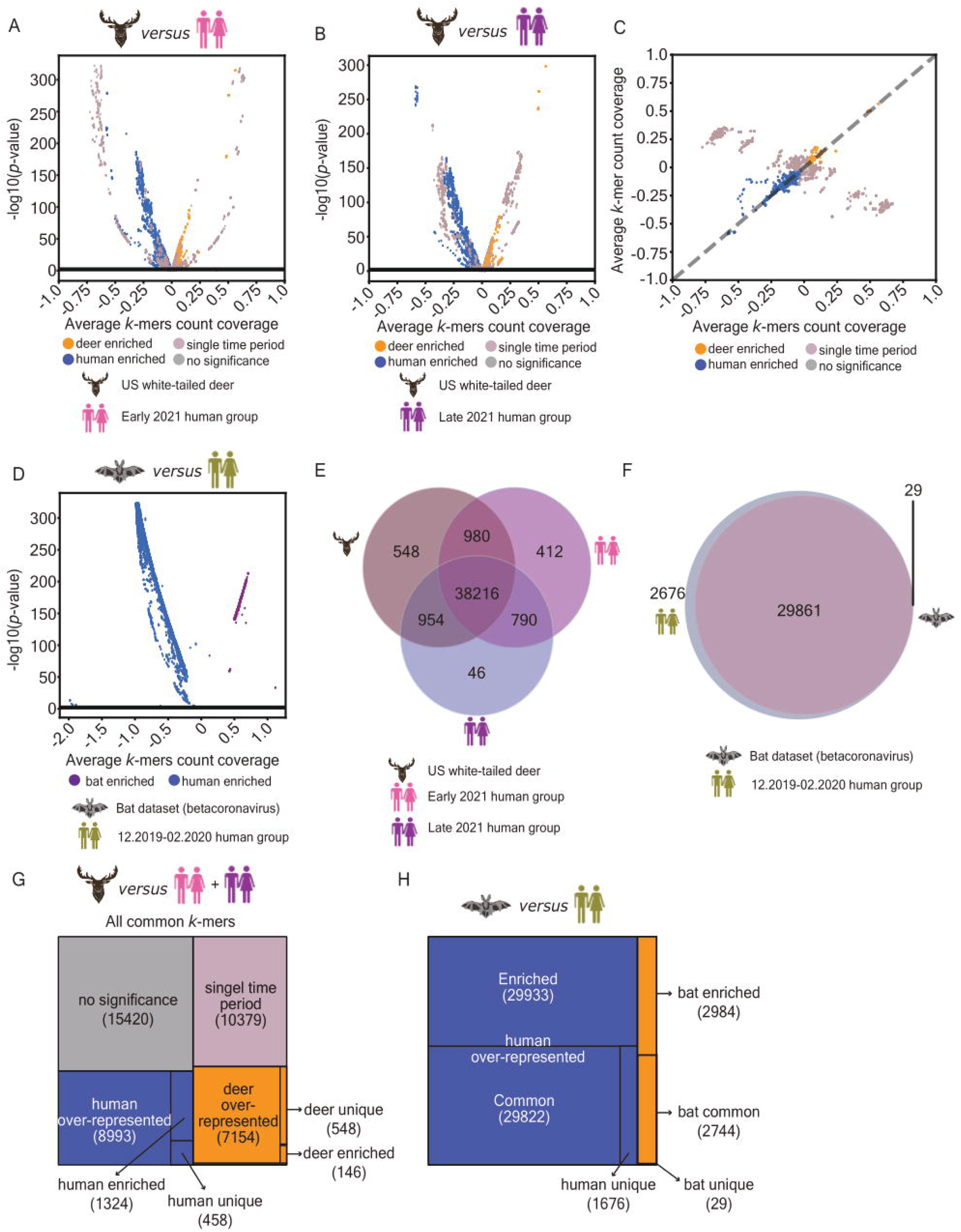
Identification of enriched *k*-mers. (**A**, **B**) Volcano plots representing *k*-mers that were significantly over-represented in deer isolates compared to isolates from the early (**A**) and late (**B**) 2021 human group, respectively. Each spot indicates one unique and over- represented *k*-mers collected after the third layer of filtering. Spots marked in orange represent *k*-mers over-represented in deer isolates; spots marked in blue represent *k*-mers over-represented in human isolates; spots marked in mauve represent *k*-mers that were over- represented in deer isolates against those isolated from either one of the human groups (namely single time period); spots marked in light gray represent the rest of the *k*-mers without significant enrichment. (**C**) Scatter plot specifying all subsets of *k*-mers that were present in both Fig. 2A and 2B, allowing the focus on over-represented *k*-mers generated by intrahost transmission within US white-tailed deer. Only *k*-mers over-represented in deer isolates against isolates in both early and late human groups were used for further analyses. Each spot indicates one unique and significantly over-represented *k*-mer. Spots marked in orange represent *k*-mers over-represented in deer isolates; spots marked in blue represent *k*- mers over-represented in human isolates; spots marked in mauve represent *k*-mers that were over-represented in deer isolates against those isolated from either one of the human groups (namely single time period); spots marked in light gray represent the rest of the *k*-mers without significant enrichment. (**D**) Volcano plot representing enriched *k*-mers significantly enriched in bat isolates compared to human isolates. Each spot indicates one unique and over-represented *k*-mers. Spots marked in purple represent *k*-mers over-represented in bat isolates, and spots marked in blue represent *k*-mers over-represented in human isolates. (**E**) Venn diagram representing the number of overlaps among a pool of common *k*-mers retrieved in deer isolates, and isolates from the early and late 2021 human groups. (**F**) Venn diagram representing the number of overlaps between a pool of common *k*-mers retrieved in bat and human isolates. (**G**) Treemap representing the hierarchical proportion of retrieved *k*- mers after manifesting the second layer of filtering in the deer dataset. Squares marked in orange represent *k*-mers over-represented in deer isolates; squares marked in blue represent *k*-mers over-represented in human isolates; a square marked in mauve represents *k*-mers that were over-represented in deer isolates against those isolated from either one of the human groups (namely single time period); a square marked in light gray represents the rest of the *k*-mers without significant enrichment. (**H**) Treemap representing the hierarchical proportion of retrieved *k*-mers after manifesting the second layer of filtering in the bat dataset. Squares marked in orange represent *k*-mers over-represented in bat isolates, and squares marked in blue represent *k*-mers over-represented in human isolates.

In summary, we obtained 41,946 common *k*-mers in the deer dataset: 7,154 are over-represented in deer isolates, 8,993 are over-represented in human isolates, 10,379 are over-represented in deer isolates, against only one of the two groups of human isolates, and for the remaining 15,420, the differences were not statistically significant (**Fig. 2A** and **2B**). In total, 32,566 common *k*-mers were found in the bat dataset: 2,744 are over-represented in bat isolates and 29,822 are over-represented in human isolates (**Fig. 2D**). After adding closely matching rare *k*-mers (**Fig. 1B** and detailed in **Materials and Methods**) and RMSE filtering, we retained 146 *k*-mers enriched in deer isolates versus 1,324 *k*-mers enriched in human isolates (**Fig. 2G**), with 2,984 *k*-mers enriched in bat isolates versus 29,933 *k-*mers enriched in human isolates (**Fig. 2H**). Intriguingly, the enriched *k*-mers count is more abundant than the total number of common *k*-mers in bat isolates after adding rare *k*-mers and RMSE filtering (**Fig. 2F**). We extrapolate that this circumstance is perhaps caused by the high diversity of SARS-CoV-2 genomic sequences isolated in bats, subsequently resulting in oodles of *k*-mers being filtered out of the pool of common *k*-mers while calculating the frequency and the average counts of *k*-mers. All *k*-mers are listed in **Tables S8** and **S9**.

Of note, 548 common *k*-mers were unique in deer isolates and 412 were unique to the isolates in the early 2021 human group, and 46 were unique to the isolates in the late 2021 human group (**Fig. 2E**); however, none of the unique *k*-mers appeared in enriched *k*-mers. We examined those unique *k*-mers and observed relatively low RMSE values, implying that these *k*-mers may not be representative of host-dependent variants. With respect to common *k-*mers in the bat dataset, 29 *k*-mers were unique in bat isolates and 2,676 in human isolates out of a pool of common *k*-mers (**Fig. 2F**). The number of *k*-mers retrieved from each layer of filtering is summarized in **Figures 2G** (deer dataset) and **2H** (bat dataset). Overall, we verified that PORT- EK is able to identify *k*-mers that are enriched after host domestication using metagenomic datasets of SARS-CoV-2 isolates collected in deer and bat datasets.

### The intrinsic properties of the SARS-CoV-2 genomic sequence are associated with host domestication

At this stage, we identified the SARS-CoV-2 genomic regions that appear to be more responsive to host domestication based on the enriched *k*-mers retrieved using PORT-EK. To better elucidate how the SARS-CoV-2 genome adapts to a new host, we further examined the viral genomic sequence at the single-nucleotide level and characterized the relationship between the intrinsic properties per site and enriched *k*- mers. We first observed that 315 and 6,140 genomic sites in the protein–coding regions overlaid with enriched *k*-mers in deer and bat datasets, respectively (**Fig. 3A** and **Fig. S1A**). Furthermore, 35 sites overlapped *k-*mers exclusively enriched in deer isolates (namely deer-exclusive sites), 7 overlapped *k*-mers preferentially enriched in deer isolates (namely deer-favorable sites), 1,909 overlapped *k*-mers exclusively enriched in human isolates (namely human-exclusive sites), 54 overlapped *k*-mers preferentially enriched in human isolates (namely human-favorable sites), and 219 overlapped the equal number of *k*-mers enriched in deer and human isolates (namely dh-comparable sites) (**Fig. 3A**). Intriguingly, deer-exclusive and deer- favorable sites were only detected in the non-protein–coding region (**Fig. 3A**); the majority of human-exclusive sites overlapped the SARS-CoV-2 *orf10* gene followed by *orf7b*, *orf8*, and the gene encoding the M protein in the deer dataset (**Fig. 3C** and **Fig. S1C**). In the case of human-favorable sites (**Fig. 3C** and **Fig. S1C**), the vast majority of sites overlaid the *orf10* genes.

**Fig. 3.**
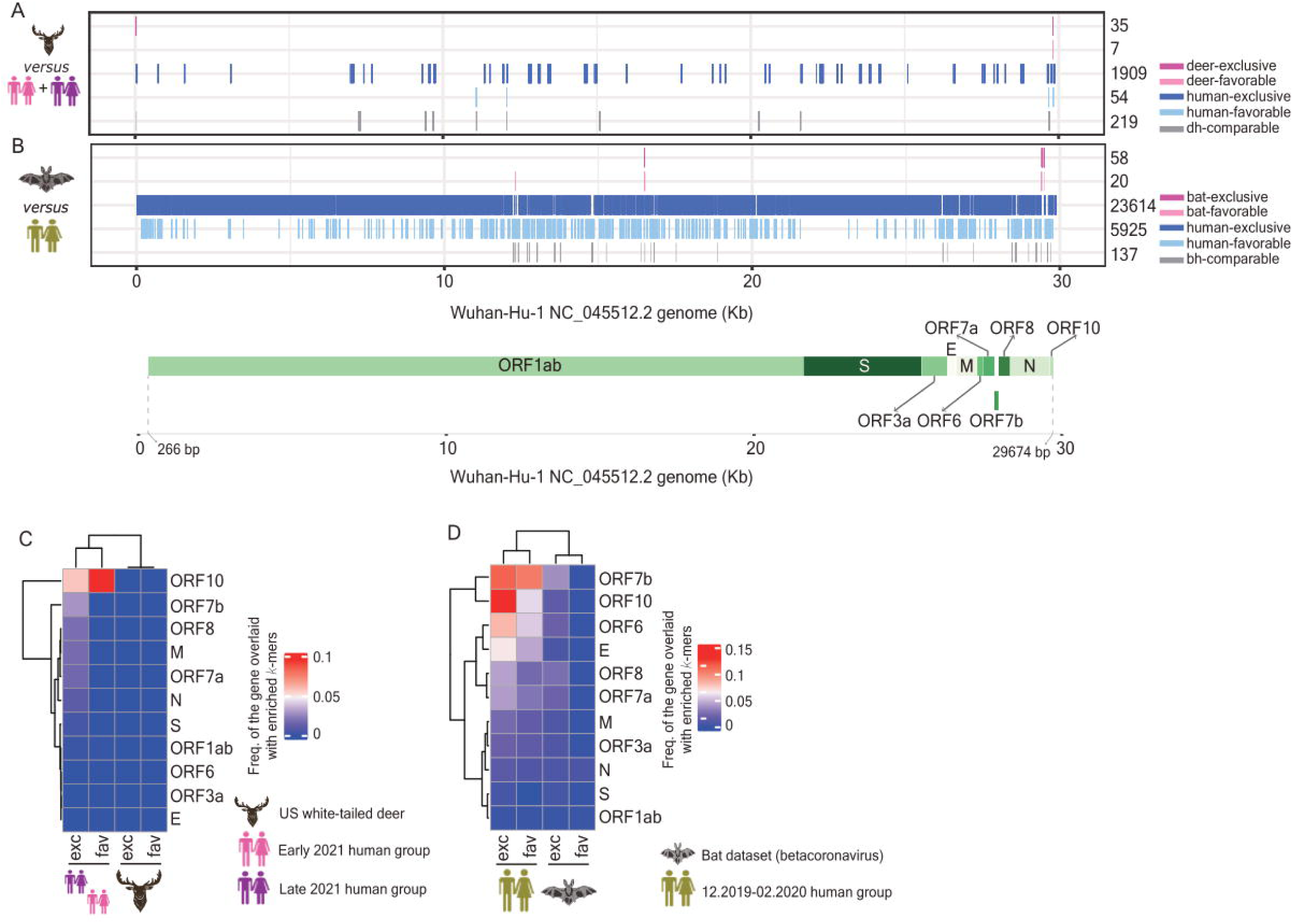
Landscapes of enriched *k*-mers throughout the whole SARS-CoV-2 genome. (**A**, **B**) Line plots visualizing the genomic loci, which present overlaps with enriched *k*-mers identified in the deer (**A**) and bat (**B**) datasets. Lines marked in dark pink represent loci overlaid with *k*- mers exclusively enriched in either deer (**A**) or bat (**B**) isolates; lines marked in light pink represent loci overlaid with *k*-mers preferentially enriched in deer (**A**) or bat (**B**) isolates compared to human isolates; lines marked in dark blue represent loci overlaid with *k*-mers exclusively enriched in human isolates in both datasets; lines marked in light blue represent loci overlaid with *k*-mers preferentially enriched in human isolates in both datasets; lines marked in grey represent the coverage of enriched *k*-mers between isolates from animal hosts (deers, panel **A**; bats; panel **B**) and humans are comparable. Gene compositions represented by a horizontal green bar were depicted on a scale of the Wuhan-Hu-1 NC_045512.2 genome. (**C**, **D**) Clustering heatmap representing the frequency of SARS-CoV-2 genes possessing enriched *k*-mers identified using the deer (**C**) and bat (**D**) datasets.

With respect to the *k*-mers enriched in the bat dataset, 58 sites overlapped *k*-mers exclusively enriched in bat isolates (namely bat-exclusive sites), 20 overlapped *k*-mers preferentially enriched in bats (namely bat- favorable sites), 23,614 overlapped *k*-mers exclusively enriched in humans (namely human-exclusive sites), 5,925 overlapped *k*-mers preferentially enriched in humans (namely human-favorable sites), and 137 overlapped an equal number of *k*-mers enriched in bats and humans (namely the bh-comparable sites) (**Fig. 3B** and **Fig. S1B**). The majority of bat-exclusive sites overlaid the *orf7b* gene followed by *orf8*, *orf7a*, and *orf6*; the majority of bat-favorable sites overlaid the gene encoding the N protein and a tiny fraction of the sites overlaid the *orf1ab* gene (**Fig. 3D**). Once again, in the bat dataset, the vast majority of human-exclusive sites overlaid the *orf10* genes followed by *orf7b*, *orf6,* and the gene encoding the E protein (**Fig. 3D** and **Fig. S1D**); in the case of human-favorable sites, the majority of the sites overlaid the *orf7b* genes followed by *orf10* and *orf6* (**Fig. 3D** and **Fig. S1D**). It is important to stress that a clear separation of two clusters based on single-nucleotide positions overlaying enriched *k*-mers was identified in animal host isolates versus human isolates (**Fig. 3C** and **3D**), signifying the differences in the intrinsic properties among SARS-CoV-2 isolates after host domestication and the feasibility of using PORT-EK for the identification of *k*-mers associated with specific host reservoirs.

### Signatures of mutations present in enriched *k*-mers at the single-nucleotide level

Mutations are one of the important mechanisms used by viruses to adapt to new environments, and whether the enriched *k*-mers identified in different host species are associated with specific signatures of mutations is now a question of interest. To tackle this question, we aligned enriched *k*-mers to the SARS-CoV-2 isolate Wuhan-Hu-1 complete genome sequence (NC_045512.2) and sought mutations emerging from individual *k*-mers. Theoretically, the coverage of each locus should be 15 because the length of a *k*-mer is set to 15; the coverage of *k*-mers above 15 indicates the presence of mutations. It is worth noting that we noticed zero (**Fig. S2A**) and 0.35% (**Fig. S2C**) of genomic loci overlaid more than 15 enriched *k*-mers in deer and bat isolates, respectively, compared to humans and zero (**Fig. S2B**) and 0.77% (**Fig. S2D**) of genomic loci overlaid more than 15 enriched *k*-mers in humans compared to deer and bats, respectively, indicating that at least one mutation has emerged at this specific locus. Although we were not able to exclude the importance of other mutations emerging in the loci covered by less than 15 enriched *k*-mers, we presume that mutations merging at loci with coverage over 15 may be more critical (here on referred to as “critical mutations”) for virus adaptation to specific hosts. Of the enriched *k*-mers identified in the deer dataset, six mutations (C7267T, C7303T, C9430T, C9679T, T15096C, and C20259T) are critical and two are not (G11083T and C29679T) (**Fig. 4A**). Among the critical mutations, C9430T and C7303T demonstrated higher frequencies than the others (**Fig. 4C**). None of the mutations emerging from the enriched *k*-mers identified in deer isolates demonstrated more than a 10% aggregated mutation frequency in *k*-mers enriched in human isolates (**Fig. S3A** and **S3B**). Except for the C29679T mutation, which emerged in the non-protein–coding region, the rest emerged in the *orf1ab* gene (**Fig. 4E**), encoding the proteins nsp3, 4, 6, 15, and RdRP. Overall, a total number of 148 mutations were identified in enriched *k*-mers in the bat dataset (**Fig. 4B** and **Table 1**); 28 mutations are critical. The occurrence of all detected conversion of mutations between unaltered and mutated nucleotides was summarized in **Table 1**. Once again, none of the mutations emerging from the enriched *k*- mers identified in the bat isolates demonstrated more than a 10% aggregated mutation frequency in *k*-mers enriched in human isolates (**Fig. S3C**).

**Fig. 4.**
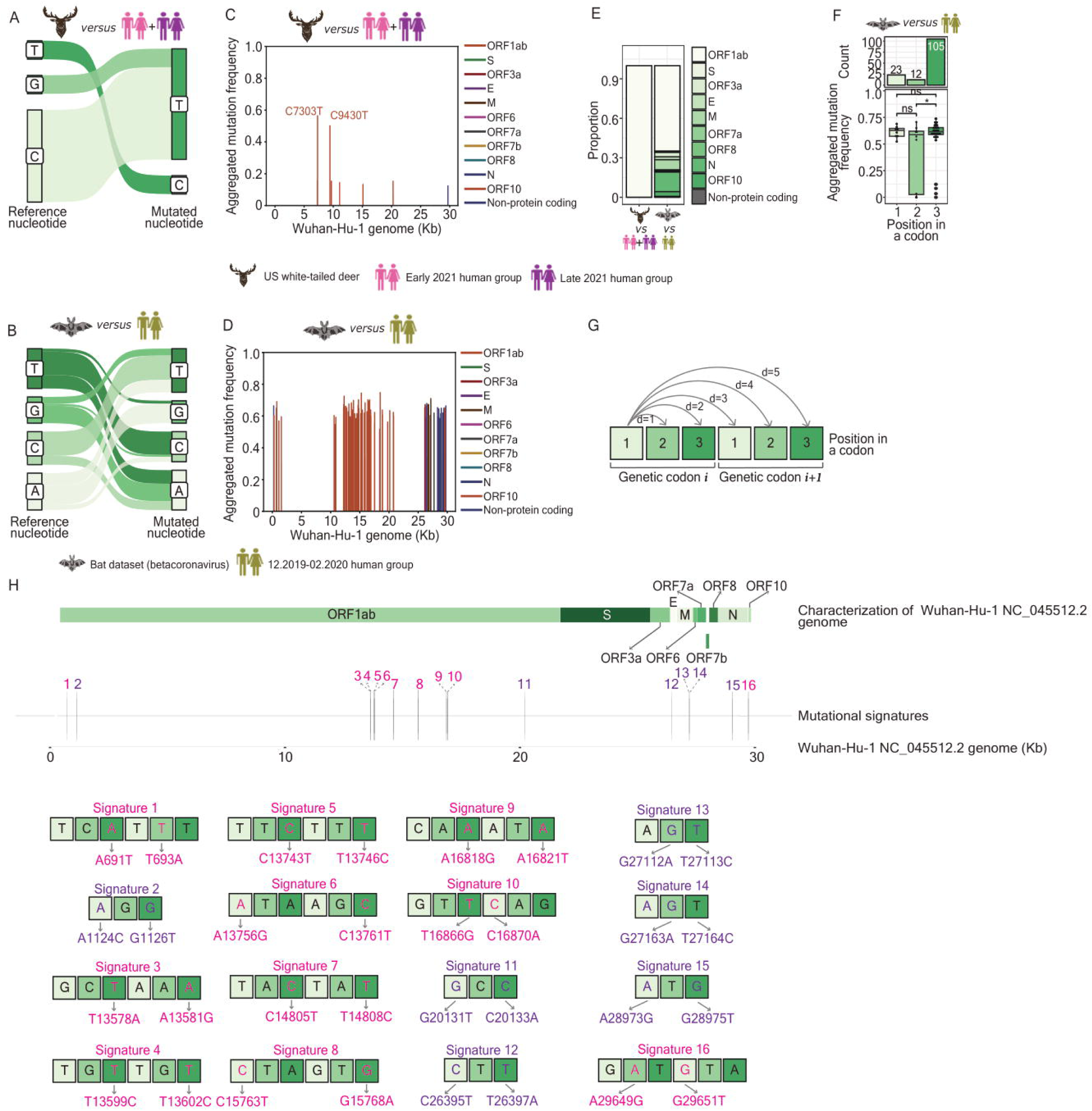
Characterization of nucleotide mutations detected from enriched *k*-mers. (**A**, **B**) Sankey plots representing the relation between reference nucleotides based on the Wuhan-Hu-1 NC_045512.2 genomic sequence and mutated nucleotides present in enriched *k*-mers identified in the deer (**A**) and bat (**B**) datasets. (**C**, **D**) Line plots representing the frequency of detected mutations per site after *k*-mer aggregation in the deer (**C**) and bat (**D**) datasets. (**E**) Stacked bar chart representing the proportion of SARS-CoV-2 genes harboring nucleotide mutations in enriched *k*-mers identified in the deer and bat datasets. (**F**) Bar chart (the upper panel) and the box plot (the lower panel) representing the total number of nucleotide mutations (the upper panel), separated by the position within a triplet of a codon encoding an amino acid and the corresponding aggregated mutation frequency (the lower panel) computed from enriched *k*-mers identified in bat dataset. Significance levels are denoted as follows: * *p* 0.05. (**G**) Schematic representation of the principle of the “signature” defined in this study: we recaptured the two mutated nucleotides with a maximum distance of 5 nt long, resulting in either a single amino acid mutation or two adjacent mutated amino acids. Such a set of a single triplet or two adjacent triplets are so- called signatures in this study. (**H**) Schematic representation of 16 signatures characterized from enriched *k*-mers identified in the bat dataset.

**Table 1.** 148 nucleotide mutations emerging from enriched *k*-mers in the bat dataset. A summary of the occurrence of the conversion of mutations between unaltered and mutated nucleotides.

| Conversion of nucleotide mutations | Occurrence |
| --- | --- |
| A to T conversion | 14 |
| A to C conversion | 6 |
| A to G conversion | 20 |
| T to A conversion | 14 |
| T to C conversion | 25 |
| T to G conversion | 4 |
| C to A conversion | 10 |
| C to T conversion | 27 |
| G to A conversion | 19 |
| G to T conversion | 7 |
| G to C conversion | 2 |

Of a total number of 148 mutations identified from the enriched *k*-mers in the bat dataset, the majority emerged from *orf1ab* (n = 91; 61.49%), followed by the gene encoding the N (n = 22; 14.86%) and M (n = 11; 7.43%) proteins (**Fig. 4E**). Although we only detected one mutation (A23403G) emerging from the gene encoding the spike protein in enriched *k*-mers, this mutation actually overlaid the viral genomic location, where the D614G mutation was identified in SARS-CoV-2 Omicron BA.2, XBB.1.5, XBB.1.16, EG.5.1, and JN.1 (**Fig. 4D**). This finding stresses the fact that readouts produced using PORT-EK are not just speculative but can also reflect natural circumstances. Intriguingly, when analyzing the composition of mutated nucleotides, we observed that for those mutations present in the coding regions, the majority of them occur at the third position (n = 105; 75%) within a genetic codon associated with a modest increase in aggregated mutation frequency compared to those at the first (n = 23; 16.4%) and the second (n = 12; 8.6%) (**Fig. 4F**), indicating that the residue at the third position within a codon is more prone to mutation and is less stable.

We continued to seek whether any viral genomic region is more responsive to mutagenesis by calculating the distance between two mutated nucleotides. We highlighted the two mutated nucleotides with a maximum distance of 5 nt, resulting in either a single amino acid mutation or two adjacent mutated amino acids (**Fig 4G** and **4H**). We termed them “signatures”. A total number of 16 signatures were observed, separated by two scenarios: (1) two nucleic acid mutations took place within the same codon, and (2) two nucleic acid mutations took place at different and adjacent codons (**Fig. 4H** and **Table 2**). Five signatures, including #2, #11, #12, #13, #14, and #15, and twelve signatures, #1, #3, #4, #5, #6, #7, #8, #9, #10, and #16 were assigned to the former and latter scenarios, respectively (**Fig. 4H** and **Table 2**). Unaltered and mutated nucleotides with respective genomic positions and scenarios were summarized in **Table 2**. Mutated nucleotides overlying non-protein–coding regions are not visualized in **Fig. 4H**. Overall, we characterized the mutational signatures present in enriched *k*-mers identified using PORT-EK.

**Table 2.**
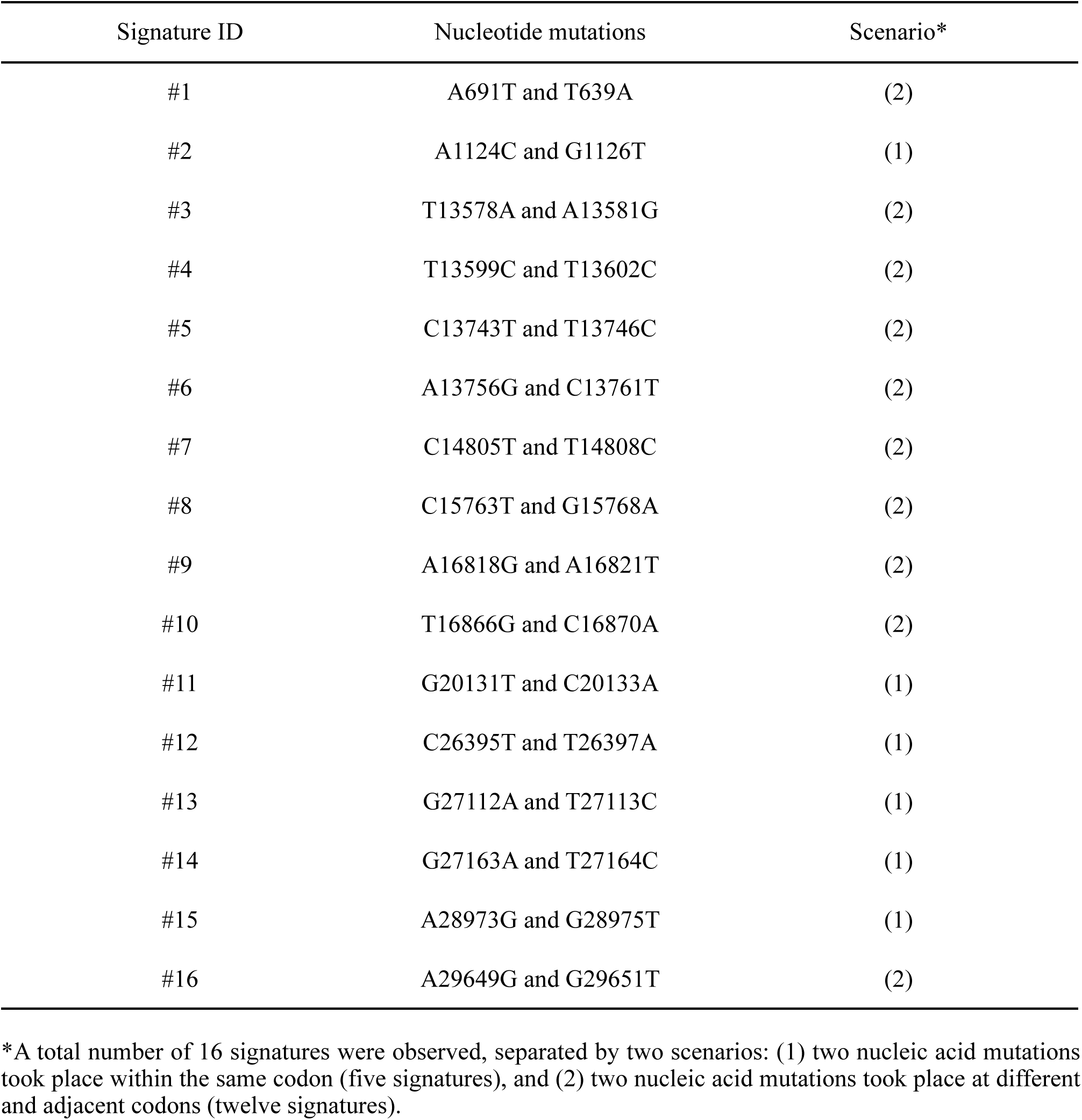
16 signatures of mutations emerging from enriched *k*-mers in the bat dataset. A summary of 16 signatures of mutations, indicating the nucleotide mutations and respective scenarios.

| Signature ID | Nucleotide mutations | Scenario* |
| --- | --- | --- |
| #1 | A691T and T639A | (2) |
| #2 | A1124C and G1126T | (1) |
| #3 | T13578A and A13581G | (2) |
| #4 | T13599C and T13602C | (2) |
| #5 | C13743T and T13746C | (2) |
| #6 | A13756G and C13761T | (2) |
| #7 | C14805T and T14808C | (2) |
| #8 | C15763T and G15768A | (2) |
| #9 | A16818G and A16821T | (2) |
| #10 | T16866G and C16870A | (2) |
| #11 | G20131T and C20133A | (1) |
| #12 | C26395T and T26397A | (1) |
| #13 | G27112A and T27113C | (1) |
| #14 | G27163A and T27164C | (1) |
| #15 | A28973G and G28975T | (1) |
| #16 | A29649G and G29651T | (2) |

\*A total number of 16 signatures were observed, separated by two scenarios: (1) two nucleic acid mutations took place within the same codon (five signatures), and (2) two nucleic acid mutations took place at different and adjacent codons (twelve signatures).

### Signatures of mutations present in enriched *k*-mers at the amino acid level

A total of seven mutations emerging from enriched *k*-mers identified in the deer dataset were present in the protein–coding regions. Six mutations resulted in synonymous mutations, including phenylalanine (Phe) to Phe (C7267T; nsp3, C9679T; nsp4, and C20259T; nsp15), isoleucine (Ile) to Ile (C7303T; nsp3 and C9430T; nsp4), and asparagine (Asn) to Asn (T15096C; RdRP) (**Fig. 5A**, **5C**, and **5E**). One mutation led to leucine (Leu) to Phe mutation (G11083T; nsp6) (**Fig. 5A**, **5C**, and **5E**). G11083T emerged in human-favorable sites; the rest emerged in dh- comparable sites.

**Fig. 5.**
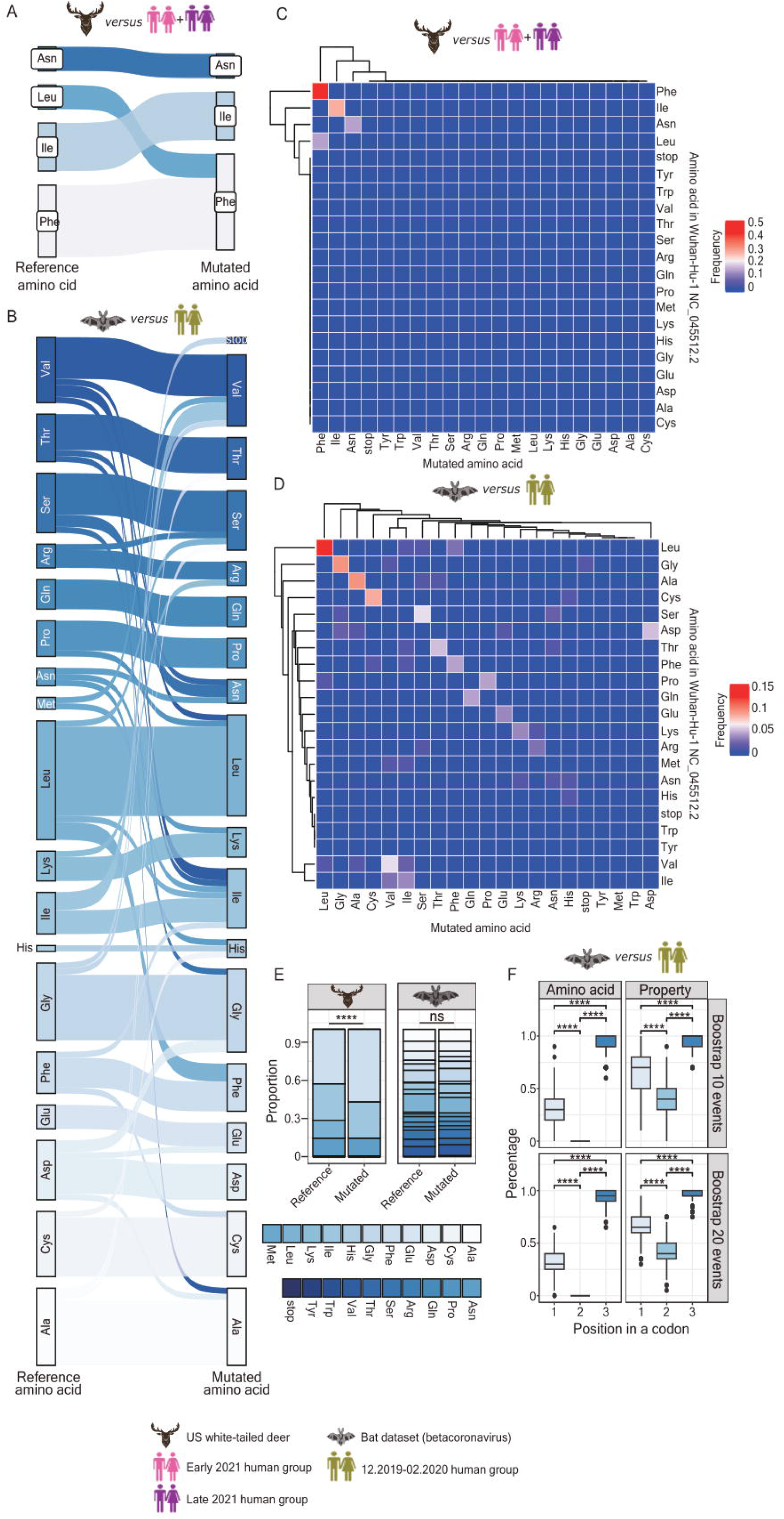
Characterization of mutated amino acids identified from enriched *k*-mers. (**A**, **B**) Sankey plots representing the relation between amino acids encoded by unaltered nucleotides based on the Wuhan-Hu-1 NC_045512.2 genomic sequence and mutated amino acids from enriched *k*-mers identified in the deer (**A**) and bat (**B**) datasets. (**C**, **D**) Clustering heatmap representing the frequency of mutated amino acids resulting from nucleotide mutations from enriched *k*-mers identified in deer (**C**) and bat (**D**) datasets. (**E**) Stacked bar charts representing the proportion between unaltered and mutated amino acids from enriched *k*- mers identified in the deer (left panel) and the bat (right panel) dataset. (**F**) Box plots representing the percentage of synonymous mutations (upper and lower panels on the left-hand side) and the percentage of mutated amino acids, harboring the identical physicochemical property to those translated in the case of unaltered codon usage (upper and lower panels on the right-hand side), separated by different codon positions. Either 10 (upper panels) or 20 (lower panels) pairs of unaltered-mutated amino acids were bootstrapped 1,000 times to obtain statistical robustness. Significance levels are denoted as follows: * *p* 0.05, ** *p* 0.01, *** *p* 0.001, **** *p* 0.0001.

With respect to mutations emerging in enriched *k*-mers identified in the bat dataset, 140 out of 148 mutations were present in the protein–coding regions. Within those 140 mutations, 75% of the mutations encode synonymous mutations (**Fig. 5B**, **5D**, and **5E**), as a diagonal was detected in the clustering heatmap shown in **Fig. 5D**. The highest mutation frequency was Leu to Leu mutation (n = 15; frequency = 0.107) (**Fig. 5B** and **5D** and **Table 3**), followed by alanine (Ala) to Ala and glycine (Gly) to Gly mutations (n = 11; frequency = 0.0785) (**Table 3**). Mutations that occurred only once were considered nonsynonymous, except histidine (His) to His and asparagine (Asn) to Asn (frequency = 0.00714); one mutation that results in a stop codon was observed (G28631T; frequency = 0.00714) (**Table 3**). We did not observe methionine (Met), tryptophan (Trp), or tyrosine (Tyr) resulting from mutations in enriched *k*-mers identified in the bat dataset (**Fig. 5E** and **Table 3**). All pairs between unaltered and mutated amino acids were summarized in **Table 3**. The coordinates of the mutated amino acids coupled with their nucleotide mutations are listed in **Supplementary Tables S10** (deer dataset) and **S11** (bat dataset).

**Table 3.** The frequency of mutated amino acids translated from nucleotide mutations emerging from enriched *k*-mers in the bat dataset. A summary of the occurrences and frequencies of 140 amino acid mutations present in the protein–coding regions.

| Amino acid mutations | Occurrence | Frequency |
| --- | --- | --- |
| Leu to Leu | 15 | 0.107 |
| Ala to Ala | 11 | 0.0785 |
| Gly to Gly | 11 | 0.0785 |
| Cys to Cys | 10 | 0.0714 |
| Ser to Ser | 7 | 0.05 |
| Val to Val | 7 | 0.05 |
| Asp to Asp | 6 | 0.0428 |
| Thr to Thr | 6 | 0.0428 |
| Phe to Phe | 5 | 0.0357 |
| Gln to Gln | 5 | 0.0357 |
| Pro to Pro | 5 | 0.0357 |
| Glu to Glu | 4 | 0.0285 |
| Ile to Ile | 4 | 0.0285 |
| Lys to Lys | 4 | 0.0285 |
| Leu to Phe | 3 | 0.0214 |
| Arg to Arg | 3 | 0.0214 |
| Ile to Val | 3 | 0.0214 |
| Asp to Gly | 2 | 0.0142 |
| Val to Ile | 2 | 0.0142 |
| Ser to Asn | 2 | 0.0142 |
| His to His | 1 | 0.00714 |
| Asn to Asn | 1 | 0.00714 |
| Asp to Ala | 1 | 0.00714 |
| Val to Ala | 1 | 0.00714 |
| Phe to Cys | 1 | 0.00714 |
| Asp to Glu | 1 | 0.00714 |
| Ser to Gly | 1 | 0.00714 |
| Cys to His | 1 | 0.00714 |
| Asn to His | 1 | 0.00714 |
| Phe to Ile | 1 | 0.00714 |
| Leu to Ile | 1 | 0.00714 |
| Met to Ile | 1 | 0.00714 |
| Thr to Ile | 1 | 0.00714 |
| Asn to Lys | 1 | 0.00714 |
| Pro to Leu | 1 | 0.00714 |
| Val to Leu | 1 | 0.00714 |
| Thr to Asn | 1 | 0.00714 |
| Lys to Arg | 1 | 0.00714 |
| Ala to Ser | 1 | 0.00714 |
| Leu to Ser | 1 | 0.00714 |
| Arg to Ser | 1 | 0.00714 |
| Ala to Thr | 1 | 0.00714 |
| Gly to Val | 1 | 0.00714 |
| Met to Val | 1 | 0.00714 |
| Gly to stop codon | 1 | 0.00714 |

In the previous section, based on nucleotide mutations emerging from enriched *k*-mers identified in the bat dataset we demonstrated that relative to the first and the second nucleotides within a genetic codon, the nucleotide at the third position faltered in stability (**Fig. 4F**). In line with this observation, we interrogated whether the mutations that take place at the third nucleotide may differ from those occurring at the first and the second position at the protein level. To tackle this question, we segregated all mutations identified in the bat dataset into three subsets based on the position of a mutated nucleotide within a genetic codon. We bootstrapped either 10 or 20 mutation events from each subset 1,000 times and examined the frequency of synonymous mutations and the physicochemical properties of mutated amino acids (**Fig. 5F**). We first observed that relative to nucleotide mutations at the first and the second position in a codon, nucleotide mutations at the third codon position frequently result in silent mutations (**Fig. 5F**). We further designated 20 amino acids to six categories based on the specificity of their physicochemical properties, including (1) amino acid residues with positively charged side chains (Arg, His, and Lys), (2) amino acid residues with negatively charged side chains (Asp and Glu), (3) amino acid residues with polar, unchanged side chains (Ser, The, Asn, and Gln), (4) amino acid residues with special side chains (Cys, Gly, and Pol), (5) amino acid residues with hydrophobic side chains in the absence of an aromatic ring (Ala, Val, Ile, Leu, and Met), and (6) amino acid residues with hydrophobic side chains in the presence of an aromatic ring (Phe, Tyr, and Trp), based on the work of Creixell et al. (2012) (*31*) with modifications. Intriguingly, we observed the superior frequency at which the physicochemical properties of amino acids translated by codons with nucleotide mutations at the third position were identical to those translated in the case of unaltered codon usage (**Fig. 5F**). Altogether, these findings suggest the possibility that codon usage bias may be involved in the process of host domestication.

### Classification and prediction

We utilized the total count of enriched *k*-mers to predict the most probable host species of a viral genomic sequence. To reduce data and model dimensions and improve performance, *k*- mers with highly linearly correlated counts were grouped, and a single *k*-mer from each group was selected. This reduced the number of *k*-mers identified in deer and bat isolates from 1,470 to 211 and from 32,917 to 489, respectively. We combined the genomic sequences of isolates from the early and late 2021 human groups to simplify the procedure of classification and prediction. We split pooled genomic sequences from isolates into training and test sets at a ratio of 0.7 to 0.3. We utilized principal component analysis (PCA) in two dimensions to reveal the discrepancy between the genomic sequences of isolates present in different hosts based on the enriched *k*-mer count (**Fig. 6A** and **6B**). The genomic sequences from bat isolates were linearly separable from those from human isolates (**Fig. 6B**), whereas sequences from deer isolates were not (**Fig. 6A**), implying that the diversity of SARS-CoV-2 genomic sequences between bat and human isolates is wider than that between deer and human isolates.

**Fig. 6.**
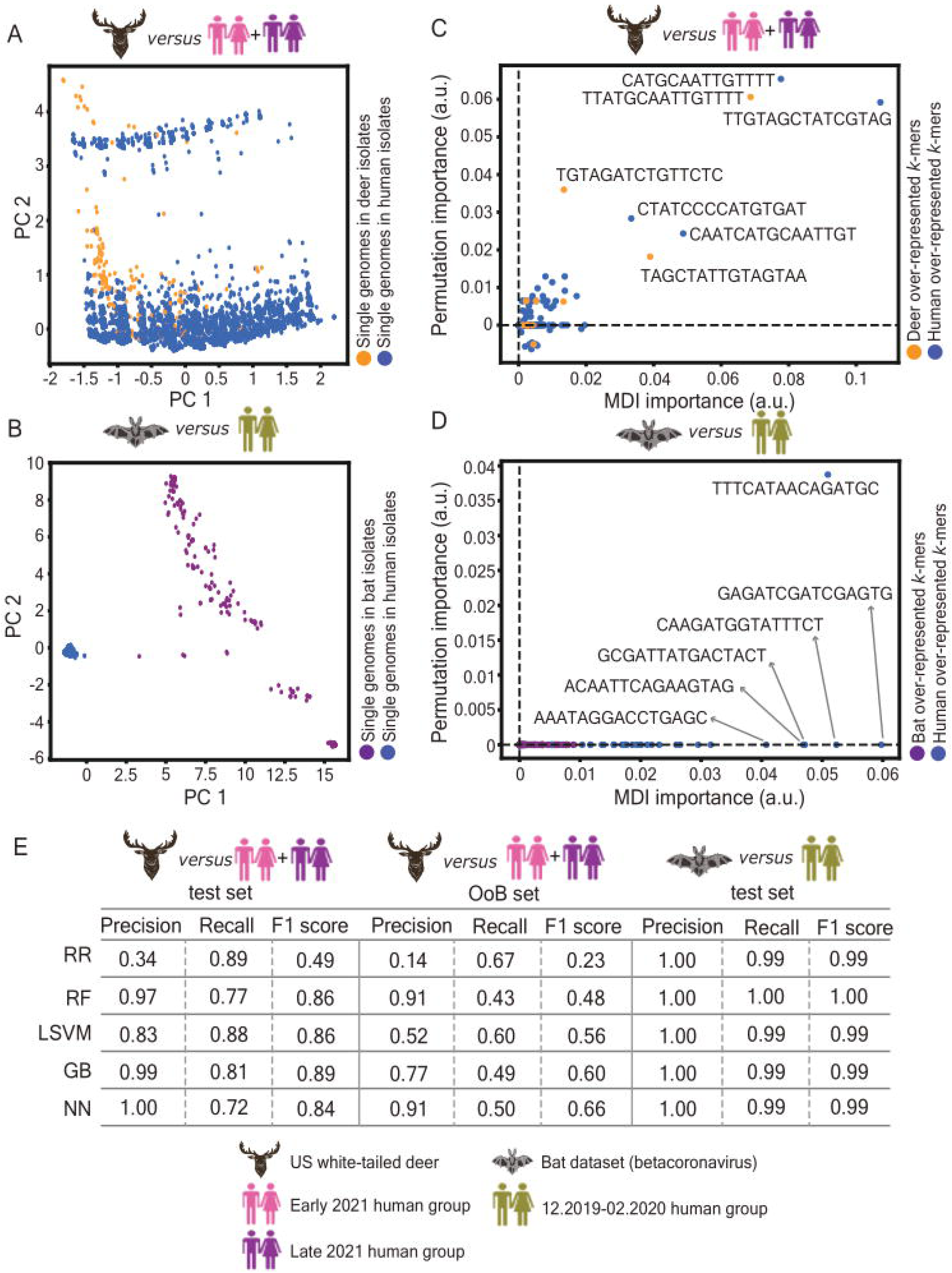
Classification and prediction of the likelihood of SARS-CoV-2 host species based on the enriched *k*-mers count. (**A**, **B**) Two dimensional principal component analysis (PCA) revealing the discrepancy of genomic sequences of isolates collected in the deer (**A**) and bat (**B**) datasets. Each dot represents a single genome of an isolate. Dots marked in orange indicate genomic sequences from deer isolates (**A**); dots marked in dark purple indicate genomic sequences from bat isolates (**B**); dots marked in blue indicate genomic sequences from human isolates (**A**, **B**). (**C**, **D**) Scatter plots unveiling the critical *k*-mer sequences for the classification of the likelihood of SARS-CoV-2 host species: deers versus humans (**C**) and bats versus humans (**D**). Based on the measures of the mean decrease of impurity (MDI) feature importance and permutation-based importance (*34*), a random forest model was constructed for classification. A total of seven (**C**) and six (**D**) enriched *k*-mers essential for the classification of the likelihood of SARS-CoV-2 host species were identified. (**E**) A table recording the record of F1 scores computed by three independent datasets: the deer dataset (on the left-hand side); the OoB dataset (middle), and the bat dataset (on the right-hand side). Five models, including ridge regression classifier (RR), random forest (RF), linear support vector machine (LSVM), gradient boosting (GB), and a dense neural network (DNN) were implemented to verify the effectiveness of each model.

We further utilized the random forest model to estimate the relative importance of *k*-mers and achieved F1 scores equal to 0.83 and 1 for the classification of *k*-mers over-represented in deer and bat isolates versus human isolates, respectively. The mean decrease in impurity (MDI) feature importance can be biased (*32*, *33*); thus, we additionally calculated permutation-based importance (*34*) to reinforce the accuracy of the classification. Although most enriched *k*-mers in deer and human isolates cluster together at low importance values, a total of seven enriched *k*-mers demonstrated the highest significance of both importance metrics (**Fig. 6C**). Intriguingly, all of these *k*-mers harbor critical mutations (C7303T, C9430T, and C9679T) that were previously identified in *k*-mers enriched in deer isolates. In contrast, the classifier failed to compute permutation-based importance in the bat dataset (**Fig. 6D**). Even though we were able to detect a total of six distinguishable enriched *k*-mers against all inputs based on MDI importance, two of them harbor critical mutations (T29758G and C14793T) identified in *k*-mers enriched in bats (**Fig. 6D**).

To ensure the robustness of our classifier, we selected the enriched *k*-mers with at least one importance measure greater than zero in an attempt to acquire the minimal dimensions in the final models and subjected them to the following models: ridge regression (RR), random forest (RF), linear support vector machine (LSVM), gradient boosting (GB), and dense neural network (DNN). Details of the model architecture are described in **Materials and Methods**. We designed F1 scores recorded from deer or bat isolates as a positive class to construct performance metrics.

Overall, the performance of the models was feasible for the classification of genomic sequences between deer and human isolates (F1 score > 0.84) and remarkable between bat and human isolates (F1 score > 0.99) (**Fig. 6E**). Among all testing classifiers, only the RR-based model failed to classify the genomic sequences between deer and human isolates (**Fig. 6E**). Confusion matrices corresponding to the individual classifiers are provided in **Figures S4A** (deer dataset) and **S4B** (bat dataset).

To further estimate whether or not our models are also applicable to other metagenomic data, we retrieved additional SARS-CoV-2 genomic sequences from deer (n = 70) and human (n = 6705) isolates within a different period of time from the metagenomic data that were previously examined in this study (detailed in **Materials and Methods**). We named this set of genomic data out of the bag (OoB) and used them to evaluate the effectiveness and feasibility of using our established models. All models, except for RR, offered acceptable performance, with DNN being the most competent (F1 score equal to 0.66) (**Fig. 6E**). The confusion matrices generated from the classifier built up using the OoB dataset are provided in **Figures S4C**.

Given that the sample size of input sequences varies in each dataset, we performed bootstrapping to generate 100 independent train–test splits and retained the DNN-based model to examine whether the varying sample sizes biased the accuracy of our models. We plotted the distribution of the F1 scores for each dataset (the deer and OoB datasets, **Fig. S5A**; the bat dataset, **Fig. S5B**). We observed, once again, the robustness of the DNN-based model for the classification of genomic sequences possessing enriched *k*-mers toward respective hosts, indicating that the DNN-based model could serve as a default choice for prediction while using PORT- EK. Altogether, these findings indicate that the enriched *k*-mer count computed by PORT-EK could be sufficient when predicting the likelihood of a host using metagenomic genomic sequence data.

## Discussion

Zoonotic viruses have caused severe disease outbreaks, ranging from sporadic cases to pandemics. Multiple factors involved in ecology, geography, human population density (*1*), alterations in human-related land use (*35*) and viruses themselves are tightly associated with zoonotic risk. As such, interrogating the mechanisms of zoonotic host jumps holds paramount importance when intervening in the spread of diseases.

Given the rapid generation of genomic datasets after the outbreak of COVID-19, appropriate machine learning-based approaches to process this massive amount of genomic information are urgently required. In this study, we establish PORT-EK with the aim of identifying enriched viral genomic regions between two metagenomes of SARS-CoV-2 isolates from different host species. PORT-EK consists of four layers of filtering steps (**Fig. 1B**), facilitating the unveiling of viral genomic regions that are enriched in one host while comparing to genomic sequences from other host isolates. Methods embedded in the filtering strategy include the calculation of frequency, the average count of *k*-mers, and RMSE. RMSE was applied as the primary performance metric, given its significance in quantifying prediction accuracy. Within the PORT-EK pipeline, we run two rounds of RMSE filtering with different purposes: the first round of RMSE performed in the third layer of filtering aims to reveal the *k*-mers that are over-represented within a specific host species, whereas the second in the fourth layer of filtering aids to diminish the background noise of filtered *k*-mers, allowing for the acquisition of the *k*-mers enriched in a specific host species. Based on the calculation of the mean *k*-mer counts RMSE, we determine that *k*-mers with a length of 15 nt are optimal for SARS-CoV-2 genomic sequences from the deer and bat isolates tested in this study. The optimal length of *k-* mers should be estimated whenever the viral genomic sequences of isolates from new hosts are used.

One fundamental limitation of high-throughput DNA sequencing is the appearance of sequencing errors (*36*, *37*). For example, it has been reported that Illumina sequencing machines produce errors at a rate of ∼ 0.1-1 × 10^−2^ per nucleotide sequenced, depending on the data-filtering scheme used (*36*, *38*). To mitigate such artifacts that may result in a mislabeling of *k*-mers using metagenomic data, at the third layer of filtering, we re-examined the similarity between every pair of sequences consisting of one rare and one common *k*-mer. When their sequences were either identical or had a mismatch with a maximum of two nucleotides, we added these rare *k*-mers to common *k*-mers, generating a pool of *k*-mers over-represented in the respective hosts (**Fig. 1B**). After this procedure, followed by a fourth layer of filtering, we observed that the pool size of enriched *k*-mers was larger than the common *k*-mers identified in the bat dataset (**Fig. 2H**); however, the opposite was true in the deer dataset (**Fig. 2G**), suggesting a wide range of sequence diversity from isolates collected in the bat dataset. Whether or not such sequence diversity correlates with the better adaptation of coronaviruses in bat reservoirs compared to other animal hosts still requires further investigation, but the observation of a few unique *k*-mers over-represented in bat isolates may imply that the majority of genomic sequences of SARS-CoV-2 isolates curated from open sources may have experienced frequent cross transmission between bat reservoirs and humans. In contrast, the identification of a more abundant number of unique *k*-mers over-represented in deer isolates (**Fig. 2E**) may lead us to assume that the circulation of these SARS-CoV-2 isolates is mainly restricted to deer (US white-tailed deer) in the same geography, rendering their genomic characterization more distinguishable from isolates from other animal hosts.

A large body of evidence has shown that the success of variants of concern is enabled by the altered intrinsic functional properties of SARS-CoV-2 and, to various degrees, subsequent changes in virus transmissibility and antigenicity (*39*). However, studies have mainly investigated the gene encoding the spike protein, and whether the properties of other SARS-CoV-2 genes are influential remains elusive. Given that enriched *k*- mers identified using PORT-EK cover the complete SARS-CoV-2 genome irrespective of the genetic boundary, we thus attempted to align individual enriched *k*-mers with the viral genome per single nucleotides and observed a distinct pattern of the genomic loci overlapped enriched *k*-mers in deer, bat, and human isolates in both deer and bat datasets (**Fig. 3A** and **Fig. S1**). In this study, we determined the optimal length of *k*-mers as being 15 nt, indicating that, theoretically, the maximum *k*-mer coverage should be 15. Notably, we observed that certain loci with *k*-mer coverage over 15 (**Fig. S2**), implying that these loci might be more correlated to the domestication of respective hosts.

Enriched *k*-mers identified using PORT-EK not only reflect quantitative differences in terms of the genomic regions between isolates from two species but also possess biological insights into the host adaptation of coronaviruses. Based on these findings, we hypothesize that perhaps the intrinsic properties of the SARS-CoV-2 genome can be reshaped during the intrahost evolutionary progress of host domestication, enabling the survival of these viruses in a new host reservoir; such intrahost selection pressure may differ among different hosts based on our findings that the loci marked by enriched *k*-mers vary between different host isolates. A clear separation of two clusters based on single-nucleotide positions overlaying enriched *k*-mers identified in animal host isolates and human isolates (**Fig. 3C** and **3D**) also strongly implies the differences in the intrinsic properties of isolates between different host species. In what way and how intrahost evolutionary constraints mechanistically reshape the intrinsic properties of a virus requires further investigation.

Furthermore, we only identified one mutation (A23403G) emerging from the enriched *k*-mer overlying the gene encoding the spike protein. Given that PORT-EK is designed to seek the enrichment of viral genomic regions based on the abundance of sequence reappearance in a dataset, we assumed that this phenomenon might be due to a high degree of genetic sequence variations present in the gene encoding the spike protein in isolates collected in the bat dataset; thus, the probability of identifying enriched genomic regions associated with the spike protein is diminished. Even so, it is remarkable that the A23403G nucleotide substitution actually results in the D614G amino acid mutation, one of the critical mutations that frequently appears in Omicron lineages (*40*, *41*), underlining the biological significance of the results generated from PORT-EK.

In this study, we further observed that nucleotide substitution at the third (wobble) position within a genetic codon is relatively flexible with a relatively higher aggregated mutation frequency relative to that measured from the other two codon positions (**Fig. 4F**), implying that, based on enriched *k*-mers identified from the metagenomes used in the bat dataset, codon bias exists, with a lack of codon stability being observed at the third position within a codon. Only a few numbers of nucleotide mutations identified in the deer dataset limit a possible verification of whether such a bias is prevalent in isolates from different host species or unique to bat isolates. A greater abundance of metagenomics data from a wide variety of host species will be required for further verification and investigation. Nevertheless, our observations lighten a potential linkage between codon stability and mutational biases resulting from, for example, APOBEC-mediated RNA editing or the role of RdRP. Additionally, the interplay between codon usage bias and host domestication will require further study.

We also observed that mutated amino acids translated from codons with nucleotide mutations at the third position tend to be synonymous (**Fig. 5F**). Relating to this, for many sets of synonymous codons in biological organisms, the base composition at the first and second codon positions are fixed and vary only at the third position (*42*). This is the so-called codon bias. Such bias has been characterized in both single-cell organisms (prokaryotes, archaea, and some fungi) (*43*) as well as different viral families (*44*). In the former case, the bias of codon usage can be attributed to isochore-dependent genome composition (GC) content, gene architecture, chromosomal locations, and endogenous gene expression (*43*, *45*). In addition, evolutionary forces and multiple molecular processes, e.g., unbiased gene conversion, mutation rates, and genetic drift, have also been discussed (*46*, *47*). In the latter case, bacteriophages, for example, their genomes are under codon-selective pressure influenced by the translational biases of their respective hosts (*48*). For most poxviruses, codon usage is close to the value predicted based on the GC content (*49*). Similar results have been obtained for vertebrate-infecting DNA viruses (*50*). In papillomavirus, codon bias was, however, attributed to the AT content (*51*). For retroviruses, strong discrimination against CpG sequences was shown to shape codon usage (*51*, *52*).

Codon usage patterns have been obtained for several members of human and non-human coronaviruses. Relative to SARS-CoV Tor2, SARS-CoV Urbani, and MERS-CoV HCoV-EMC, SARS-CoV-2 (*46*) and human coronaviruses (HCoV-OC43, HCoV-HKU1, HCoV-229E, and HCoV-NL63) display AU-rich bias in terms of codon usage (*53*, *54*), indicating that codon usage bias can vary among coronavirus isolates from different host reservoirs and be shaped by mutation pressure and evolutionary constraints. The same result was reported by Gu et al. (2004) (*55*), in that a slight bias of synonymous codon usage is present in the SARS-CoV genes with a preference for A-ended or U-ended codons. Analogous to this finding, we observed a higher frequency of mutated nucleotides identified on enriched *k*-mers in the bat dataset in terms of A and T (A, 29.2%; T, 32.7%) (**Fig. 4B**).

Among the 20 standard amino acids, Leu, Ser, and Arg, which can be encoded by six triplets of the genetic codons, are those with the widest range of codon usage. Our observation also highlights the superior frequency of the Leu-to-Leu mutation emerging in enriched *k*-mers identified in the bat dataset (**Fig. 5D**). This finding could be supported by the findings of Hou (2020) (*54*), indicating that Leu is one of the most frequently used amino acids in all human coronaviruses analyzed. Codon usage bias can also affect messenger RNA (mRNA) stability, subsequently influencing gene expression and modulating the translation of protein synthesis (*56*, *57*). Whether the destabilization of the third residue also influences the RNA and protein synthesis of SARS-CoV-2 remains elusive and is in urgent need of study.

Earlier research showed that amino acid substitutions occur less frequently between amino acids with very different physicochemical properties (*58*) based on various codon substitution models (*59-64*). Notably, in this study, we also observed that a vast majority of amino acids translated from the codons with a mutation at the third position possess physicochemical properties identical to those encoded by an unaltered genetic codon (**Fig. 5F**). Such a correlation may imply that, in addition to the position of the genetic codon, the physicochemical properties of the amino acid could also be reshaped by evolutionary selection pressure. Further studies focusing on how these two parameters, genetic codon position and the physicochemical properties of an amino acid, concomitantly or independently, govern alterations in coronavirus tropism associated with different host reservoirs, especially in the context of determining synonymous codon usage, would facilitate predicting the evolutionary trajectories of coronaviruses across different host species.

Nevertheless, viruses show considerable variation in the selectivity with which they infect host species, and the features that govern host adaptation vary between different viruses. In this study, we presented the analytical pipeline of PORT-EK, which was used to compare metagenomes from two hosts and identify the genomic regions that are more quantitatively dominant in one host than another. One of the significant advances of this pipeline is that it allows for the identification of silent mutations emerging in viral genomic regions over-represented in respective hosts because PORT-EK enables direct metagenomes at the nucleotide level. How such synonymous codon usage bias subsequently influences host adaptation and the cross-species transmission of coronaviruses will require further investigation. In addition, knowledge of the characterization of different codon usage patterns in SARS-CoV-2 variants will have potential value for developing coronavirus vaccines. Historical outbreaks signify that the further characterization of the animal hosts implicated in the spillover or spillback events of coronaviruses would be beneficial for current pandemic management and the prevention of imminent outbreaks. We envision that PORT-EK will constructively support the investigation of massive metagenomes with a resolution of single nucleotides.

### Limitations of this study

One of the limitations is that we cannot ascertain whether enriched *k*-mers identified using PORT-EK cover all dominant genomic regions resulting from host domestication. Undiscovered *k*-mers may be due to sequences of isolates that are either not yet characterized or not included in the present dataset. This issue is also referred to that the sample size in some species, e.g. the genomic sequences in deer isolates, is small in this study. The fewer sequences are available, the less significant the *p*-value is, rendering it difficult to identify enriched *k*-mers. Furthermore, in this study, we did not investigate and compare enriched *k*-mers identified from multiple periods, rendering it difficult to track their evolutionary trajectories in circulation within a host. Indeed, based on PORT-EK, we identified enriched *k*-mers with a statistical test that assumes independence; the frequencies of the *k*-mers are however not independent in the real world: viruses evolve biologically so a mutation that appears early will remain in all the descendants until it mutates again. Further advanced statistical methods, e.g. phylogenetic regression will be required to perform statistics on samples of biological sequences.

It is known that viruses often possess variations within an individual host and exist as a population of variants; several factors, including the discrepancy of mutation profiles between variants within a host and consensus single-nucleotide polymorphisms observed in the genomic sequences among SARS-CoV-2 variants, strengthen the importance of evolutionary intrahost constraints that reshape the intrinsic properties of the SARS-CoV-2 genome (*65*, *66*). This question is crucial, and when properly addressed, may facilitate a deeper understanding of viral–host adaptation and cross-species transmission driven by evolutionary intrahost constraints. Here it is also important to note that theoretical findings shown in this work should also be further verified using appropriate experimental models.

In the last four years, various *k*-mer-based algorithms have been developed and utilized for the detection of SARS-CoV-2 variants of concern, classified by specific geographical regions during a certain period, aligning with the principles of novelty detection (*20*, *29*, *67-76*) and for the identification of mutations emerging during intrahost circulation (*77*). Consistently, PORT-EK is capable of processing, with a wide range of metagenomic sequencing datasets and seeking genomic regions over-represented in respective hosts. At present, only a small fraction of viral diversity circulating in wild and domestic vertebrates has been characterized. Thus, we envision that PORT-EK is also applicable to other zoonotic viruses. In this circumstance, we recommend the unique parameterization of PORT-EK to optimize its effectiveness toward different zoonotic viruses.

Technically, in the PORT-EK pipeline, our filtering strategy for the selection of enriched *k*-mers is based on the frequency of *k*-mers, the difference in the average counts of *k*-mers between two species, and RMSE, and other filtering methods (*78*), such as minimizers and convolutional neural networks (CNNs), will also be implemented to reinforce the accuracy of PORT-EK. In addition, it is important to stress that when evaluating the robustness of PORT-EK, we noticed that some enriched *k*-mars cannot be aligned to the reference genome. We assume that it may be due to (1) the choice of the mapping algorithm or (2) the fact that the reference genome chosen cannot cover all SARS-CoV-2 variants. In the future, the selection of multiple reference genomes and mapping algorithms should also be considered to reinforce the completeness of the final readouts.

Finally, we propose that the usage of PORT-EK should be coupled with other genomic approaches, such as RNA-seq, single-cell RNA-seq, and Oxford Nanopore long-read sequencing, to augment the ramifications of the identified *k*-mers. Nevertheless, we offer a perspective lens on viewing potential SARS-CoV-2 genetic markers for the classification of metagenomic viral sequences resulting from host domestication using a *k*- mer-based analytical pipeline, PORT-EK.

## Materials and Methods

### Source of Data

In this study, we captured metagenomic datasets of SARS-CoV-2 isolates from US white-tailed deer and bats and compared viral genomic sequences with those retrieved from humans. All sequences were deposited either from the GISAID Initiative database (https://gisaid.org/) (*79*) or the NCBI Virus database (https://www.ncbi.nlm.nih.gov/labs/virus/vssi/<u>#/</u>) (*80*). We generated three metagenomic datasets, namely the deer, bat, and out of the bag (OoB) datasets. Each dataset consists of genomic sequences of SARS-CoV-2 isolates from two different host species, allowing the comparison of the enrichment of *k*-mers between each other.

In the deer dataset, 336 genomic sequences of SARS-CoV-2 variants collected from North America white-tailed deer between October and December 2021 were utilized (GISAID EPI_SET_240422va, available at https://doi.org/10.55876/gis8.240422va, **Supplementary Table S1**). We compared sequences isolated from deer with the sequences from human isolates collected at two different collection periods: 21,906 sequences collected in April 2021 (namely the early 2021 human group in Main Text, GISAID EPI_SET_240422rw, available at https://doi.org/10.55876/gis8.240422rw, **Supplementary Table S2**) and 11,525 sequences collected in November 2021 (namely the late 2021 human groups, GISAID EPI_SET_240422qc, available at https://doi.org/10.55876/gis8.240422qc, **Supplementary Table S3**).

With respect to out of the bag (OoB) dataset, 70 additional SARS-CoV-2 sequences collected between October and December 2021 in US white-tailed deer were used (GISAID EPI_SET_240422oy, available at https://doi.org/10.55876/gis8.240422oy, **Supplementary Table S4**) and 6,705 sequences from human isolates collected in January 2022 were used (GISAID EPI_SET_244022xu, available at https://doi.org/10.55876/gis8.240422xu, **Supplementary Table S5**).

In the bat dataset, we retrieved a total of 263 available betacoronavirus sequences isolated from bats in the NCBI Virus database (**Supplementary Table S6**) and 2081 sequences collected between December 2019 and February 2020 (GISAID EPI_SET_240422qm, available at https://doi.org/10.55876/gis8.240422qm, **Supplementary Table S7**) for analyses.

### Model overview, requirements, and parameters

PORT-EK is a pipeline that captures sequences from two different metagenomic datasets and identifies over-represented *k-*mers in respective datasets. In this study, we further utilized the count of enriched *k-*mers as the predictor variable to classify the likelihood of respective hosts, in which *k-*mers are over-represented. This pipeline is written in Python 3.11, using: biopython 1.81, numpy 1.26.2, pandas 2.1.4, seaborn 0.13.2, matplotlib 3.8.2, networkx 3.2.1, scipy 1.11.4, keras 2.15.0 and scikit-learn 1.3.2. It also requires the support of Jupyter notebooks to execute several critical steps involved in PORT-EK, including *k*-mer filtering, statistics, and identification of enriched *k*-mers coupled with mutations.

While using PORT-EK, input files should be in the fasta format with headers either in GISAID or NCBI Virus formats. All sequences in one input file correspond to the same host; multiple input files can be pooled for the same host if required. PORT-EK allows the comparison of two independent sources of metagenomes. In this study, each dataset consists of metagenomes isolated from two different host species. Of note, a set of tunable parameters includes the *k-*mer length *k*, conservation threshold *c*, allowed rare *k*-mer mismatches *m*, minimum root mean square error *min_RMSE_*, allowed mapping mismatches *m_map_* , and allowed mapping offset *l_map_* are recommended to be adjusted whenever a new metagenomic dataset is applied. The details of the parameters and their settings are discussed later in the following section. Recorded running times and peak memory usage for the data sets used in this study are presented in **Supplementary Table S12.**

### *K*-mer extraction, count matrix, and descriptive statistics used in PORT-EK

We first extracted *k*-mers of the length *k* with overlapping sequences using a sliding window moving every nucleotide and placing them in the same dictionary. *K*-mer sequences and the first nucleotide position were used as indices for constructing this position matrix. A single ID was assigned to each dictionary. Host species were denoted to corresponding dictionaries of *k*-mers. Labeled indices were saved as individual files to alleviate RAM usage.

Next, the indices were read to construct a matrix containing the count of every *k*-mer in every dictionary. Inside a matrix, rows indicate the *k*-mer sequence, and columns indicate the dictionaries. *K*-mers with a poly(A) sequence were discarded. We further measured the appearance of *k*-mers, representing the “frequency” for each host. Frequency was calculated based on the ratio between the total number of *k*-mers in one dictionary and the total number of dictionaries in the host. This is the second filter, namely the rarity filter in PORT-EK (**Fig. 1B**). Based on the assumption that *k*-mers containing meaningful information should not be very rare within the hosts (*29*), only *k*-mers that are at least *c* % conserved to the respective host were retained for further analyses. Lower settings of *c* allow the pipeline to capture rarer variants that may still be meaningful at the cost of increased computation time, memory usage, and difficulty of interpretation; the reverse is true for higher settings. We found that *c* of 1% is sufficient for the deer and OoB datasets, whereas *c* of 50% is required for the bat dataset. We termed the *k-*mers that pass the rarity filter the common *k*-mers, whereas those that do not were named the rare *k*-mers.

We further computed the following three statistics on common *k*-mers:

1. the average count of each *k*-mer for two individual hosts, of a particular human host group. *n_nh_* and *n_hi_*, where *n_hi_* indicates the identifier
2. the difference in the average count of each *k*-mer between two hosts:
3. the root mean square error (RMSE) of said changes:

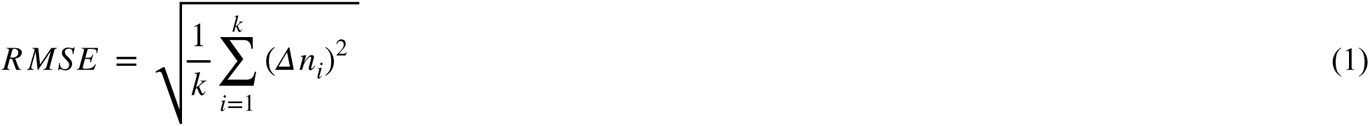

where *k* is the total number of human groups.

The difference in the average count of each *k*-mer was used throughout the whole pipeline as metrics of enrichment. In this work, the positive value denotes the average count of each *k*-mer enriched from animal hosts (deer and bats); whereas, the negative value denotes the enrichment related to humans. RMSE was used for an unsigned measure of the degree of enrichment. In other words, the larger the RMSE is, the more pronounced the difference between two comparing hosts is. Finally, the statistical significance of *Δn_i_* tested using Fisher’s exact test. The differences with *p*-values less than 0.01 were deemed significant.

### Determination of the optimal *k* value

We repeated the above statistics to determine the optimal *k* was of *k*-mers. In brief, to compare the specificity of *k*-mers with the different lengths *k,* the *n_nh_* and *n_hi_* statistics were normalized by the average count of all *k*-mers in genomic sequences of isolates within a dataset (*n_t_*), and the normalized equivalents of *Δn_i_* and *R MSE*, *Δn_i norm_*and *R MSE_norm_* , were calculated.

We further compared the *R MSE_norm_* of *k*-mers in all *k*-mers of length *k* from the whole data set (**Fig. 1C** and **1D**). In this study, we compared *k* values of 5, 7, 9, 11, 15, 19 and 25. We observed that the 99^th^ percentile *R MSE_norm_* shows an increase while the value *k* increases. A plateau appears when the *k* value is bigger than 11. Additionally, we used a fraction of *k*-mers that pass the rarity filter, *f_c_*, as a measurement of computational efficiency. The lower the required. *f_c_* is, the higher demand on computing time and memory is

### Over-represented and enriched *k*-mer identification

Over-represented *k*-mers were retrieved from a pool of common *k*-mers (**Fig. 1B**) based on three mentioned statistical strategies (the third layer of filtering). Only *k*-mers with *p*-values less than 0.01 are retained. The parameter *Δn_i_* was used to determine in which species *k*-mers are enriched. In this study, a positive value of *Δn_i_* indicates that *k*-mers are enriched in isolates from deer or bats, whereas a negative value indicates *k*-mers are enriched in human isolates. It is important to note that, with respect to *k*-mers over-represented in deer isolates, we only retain the ones that were over- represented against both early and late human 2021 groups. *K*-mers with *p*-values greater than 0.01 were discarded.

At this step, we re-examined the sequence similarity between rare and over-represented *k*-mers (**Fig. 1B**). Rare *k-*mers with the superior sequence similarity (two-nucleotides mismatches, the parameter *m*, allowed) to any of common *k*-mers were added to a pool of over-represented *k*-mers. This step is however optional in the PORT-EK pipeline. We constructed a graphic network, in which nodes represent a pool of the over- represented *k*-mers and rejoining rare *k*-mers, and edges represent the similarity between every pair of an over-represented *k*-mer and a rejoining rare *k*-mer, allowing the visualization of the distribution of both subsets of the *k*-mers.

At the final step of the PORT-EK pipeline, we applied, once again, the statistical tools implemented in the third layer of filtering to those rejoining rare *k*-mers. We passed all over-represented and rejoining rare *k*-mers to the fourth RMSE filter in order to retrieve enriched *k*-mers as a final output (**Fig. 1B**). *K-*mers that have an RMSE less than *min_RMSE_* were discarded. Of note, the parameter *min_RMSE_*is adjustable in the range between 0 and 1, allowing fine-tuning of the sensitivity of PORT-EK. As the value *min_RMSE_* is approaching 1, the strength of the fourth filter is stricter. In this study, the default setting *min_RMSE_* is equal to 0.1. The rest of the *k*-mers are considered to manifest significant enrichment and are designated to corresponding hosts. The enriched *k*-mer matrix was then transposed, as IDs were listed in rows and *k*-mers in columns. Host species were assigned to enriched *k*-mers shown in columns. Column labeling the host of the viral samples was added. Host species labels were numerical and binary: deer or bats are labeled as 1; humans as 0.

### Enriched *k-*mers mapping

In this study, we established a tailored algorithm using Python regular expression library, RegEx, enabling the mapping of the enriched *k*-mers throughout the reference genome, severe acute respiratory syndrome coronavirus 2 isolate Wuhan-Hu-1, complete genome NC_045512.2, curated in the National Center for Biotechnology Information (NCBI) database (<u>https://</u> www.ncbi.nlm.nih.gov/nuccore/1798174254). Only *k*-mers that are present one time in individual viral genomic sequences were used for mapping to minimize the probability of the presence of ambiguous *k*-mers. The genomic position corresponding to the first nucleotide from each selected *k-*mer was used as *k*-mer indices to form a matrix. In principle, 15 nt substrings in sliding windows consecutively shifting per nucleotide were read through over the whole reference genome. Every substring was compared with mentioned *k*-mer sequences with up to *i* mismatches, with *i* increasing from 0 to *m_map_* , the maximum number of mismatches.

To align enriched *k*-mers throughout the SARS-CoV-2 genome per site, we sought the lowest number of mismatches followed by the lowest starting position difference. If no substrings were found with less than *m_map_* mismatches or less than mapping offset *l_map_*, the starting position difference, and no match were returned. By default, *m_map_*, it is set to the same number as *m* denoted as the number of mismatches, at the step of the identification of rare *k*-mers, and *l_map_* is set to 1000 (both parameters are tunable). In this study, we set *m_map_* equal to 2 and *l_map_* equal to 1000 as default. Based on the collection of substrings, we could depict the viral genomic positions overlaid with enriched *k*-mers at a single-nucleotide level (**Fig. 3A**).

### Calculation of the enrichment of enriched *k*-mers overlaid with SARS-CoV-2 genes

The enrichment of enriched *k-*mers exclusively enriched in animal or human isolates at individual SARS-CoV-2 genes was calculated by Equation (1).

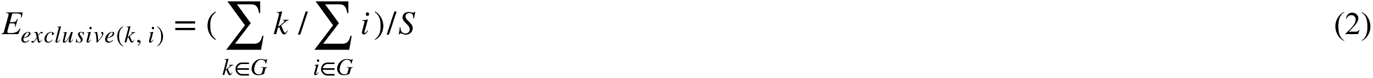

where *E_exclusive_*is the enrichment of a total number of enriched *k-*mers *k* covering given SARS-CoV-2 viral genomic loci normalized by the size of a given SARS-CoV-2 gene .

The enrichment of enriched *k-*mers preferentially enriched in either animal or human isolates at individual SARS-CoV-2 genes was calculated by Equation (2).

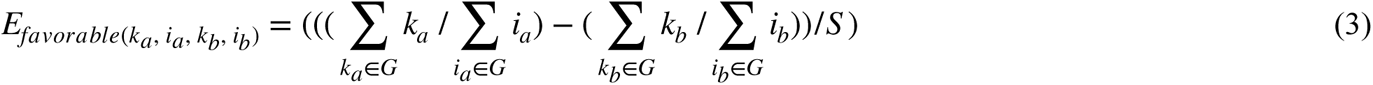

where *E_favorable_*is the enrichment of a total number of enriched *k*-mers *k_a_*, which cover given SARS-CoV-2 viral genomic loci *i_a_* in one species that dominate a total number of the enriched 15 nt *k*-mers *k_b_*, which cover given SARS-CoV-2 viral genomic loci *i_b_* in another species followed by a normalization by the size of a given SARS-CoV-2 gene . Of note, given that the total number of *k*-mers retrieved in the comparable group was identical in both species, this group was thus excluded from this analysis. The calculation of the enrichment is visualized in **Fig. S1C** and **S1D**. The genomic positions of SARS-CoV-2 protein–coding genes, including the *orf1ab* gene (266 bp -21555 bp), the gene encoding the S protein (21563 bp - 25384 bp), the *orf3a* gene (25393 bp - 26220 bp), the gene encoding the E protein (26245 bp - 26472 bp), the gene encoding the M protein (26523 bp - 27191 bp), the *orf6* gene (27202 bp - 27387 bp), the *orf7*a gene (27394 bp - 27759 bp), the *orf7b* gene (27756 bp - 27887 bp), the *orf8* gene (27894 bp - 28259 bp), the gene encoding the N protein (28274 bp - 29533 bp), the *orf10* gene (29558 bp - 29674 bp), were recorded from NCBI Reference Sequence: NC_045512.2 - Severe acute respiratory syndrome coronavirus 2 isolate Wuhan-Hu-1, complete genome.

### Calculation of the coverage of enriched *k*-mers over the SARS-CoV-2 genome

Once the single-nucleotide positions overlaid by enriched *k*-mers were characterized, we calculated the *k*-mers coverage of each genomic locus based on the total number of enriched *k*-mers overlaid. For every locus, based on the competition of the sum of enriched *k*-mers, separated by corresponding hosts, we categorized the tropism of each locus into three groups: exclusive, favorable, and comparable sites (**Fig. 3A** and **Fig. S2**). Exclusive sites were referred to as the scenario that all *k*-mers are enriched in the same host; favorable sites were referred to as the scenario that the total number of enriched *k*-mers were superior in one host species than another within the same dataset; comparable sites were referred to as the total number of enriched k-mers were equal between two host species within the same dataset (**Fig. 3A**).

### Identification of mutations on enriched *k*-mers

At the enriched *k*-mers alignment step, nucleotides that differ from those in the reference sequence, SARS-CoV-2 Wuhan-Hu-1NC_045512.2, were marked as mutations. Identical mutations present on multiple *k*-mers were aggregated. Only mutations that are supported by mappings of at least *FLOOR*(*k* /2) *k*-mers were retained. The aggregated mutation frequency was calculated using the total number of isolate sequences, in which at least one *k*-mer harboring a mutation was divided by the total number of isolate sequences in the same animal host. The highest RMSE of *k*-mers harboring mutations was recorded (**Fig. 4C** and **4D**). In parallel, we computed the aggregated mutation frequency using the total number of isolate sequences, in which at least one *k*-mer harboring a mutation divided by the total number of isolate sequences in the contrast host, humans, within the same dataset (**Fig. S3A**-**S3C**), enabling the visualization of the aggregated mutation frequency throughout the reference genome.

As a final output, we determined 8 and 148 nucleotide mutations on enriched *k*-mers identified in the deer and bat datasets, respectively. Positions of mutated nucleotides within a genetic codon and respective amino acids encoded were manually verified using the UCSC Genome Browser on SARS-CoV-2 Jan. 2020 ( N C _ 0 4 5 5 1 2 . 2 ) ( w u h C o r 1 ) ( https://genome.ucsc.edu/cgi-bin/hgTracks?db=wuhCor1&lastVirtModeType=default&lastVirtModeExtraState=&virtModeType=default&virtMode=0&nonVirtPosition=&position=NC_045512v2%3A1%2D29903&hgsid=2236086162_7owbHRSgRbADlT8C9GSdZ4z6bwp5).

### Data visualization

#### Volcano plot representation of over-represented *k*-mers

Volcano plots tailed for the representation of over-represented *k*-mers were generated using Python 3.11.6. In this plot, the x-axis represents *Δn_i_* and the y-axis represents -log_10_(*p*-value). An additional scatter plot (**Fig. 2C**) transformed from the volcano plots in the deer dataset was generated to highlight *k*-mers over-represented in deer isolates in contrast to isolates from both early and late 2021 human groups.

#### Line plot representation of SARS-CoV-2 genomic loci overlaid with enriched *k*-mers

The line plots representing the landscape of the SARS-CoV-2 genomic sequence per site, in which nucleotides overlay with enriched *k*-mers (**Fig. 3A** and **3B**) were created using the default R build-in function `geom_segment()`.

#### Clustering heatmap

The cluster heatmaps representing the frequency of the SARS-CoV-2 genes harboring enriched *k*-mers identified in deer (**Fig. 3C**) and bat (**Fig. 3D**) datasets as well as the mutation frequency between reference and mutated amino acids in deer (**Fig. 5C**) and bat (**Fig. 5D**) datasets were plotted using the R package ComplexHeatmap (*81*, *82*) with default options.

#### Sankey plot representation of the relation between reference nucleotides/amino acids and mutated nucleotides/amino acids

Sankey plots representing the transition from reference nucleotides or amino acids to mutated nucleotides or amino acids present in enriched *k*-mers identified in the deer (mutated nucleotides, **Fig. 4A**; mutated amino acids, **Fig. 5A**) and bat datasets (mutated nucleotides, **Fig. 4B**; mutated amino acids, **Fig. 5B**) were plotted using the R package ggSankey (https://github.com/davidsjoberg/ggsankey).

#### Prediction of the likelihood of the host

We utilized the counts of enriched *k*-mers as the predictor variable for the prediction of the likelihood of the host. In principle, we first calculated a correlation matrix of *k*-mer using pandas (*83*) `corr()` function and constructed a graph, in which nodes representing each enriched *k*- mer and edges connecting nodes representing *k*-mers with a correlation coefficient of more than 0.9 (including loops). The connected components shown in this graph represent the groups of *k*-mers with a high correlation of enriched *k*-mer counts. Based on enriched *k*-mer counts, we were able to calculate the absolute value of the correlation coefficient for each *k*-mer associated with the respective host and set it as a node attribute *y_corr* (**Fig. S6**). Finally, *k*-mers (illustrated as nodes in the graph) coupled with the largest *y_corr* from individual connected components in a graph were selected as the so-called linearly independent *k*-mer set.

The linearly independent *k*-mer set was split into training and test sets at a ratio of 0.7 to 0.3, with stratification associated with the respective host. The importance of particular *k*-mers for classification was calculated based on a mean decrease in impurity importance (MDI) and permutation importance (*84*) using scikit-learn’s RandomForestClassifier with balanced class_weight; remaining hyperparameters at default values. F1 score obtained from linearly independent *k*-mers enriched in deer and bats were set as positive values, serving as a metric. *K*-mers with the values of importance equal to zero or negative values of importance were removed from the dataset, and were, once again, split into training and test sets at the same mentioned ratio. Of note, given such a prevalent imbalance of the sample size of genomic sequences retrieved from animal species versus humans (*85*), rendering the measure of ROC AUC (Area under the ROC Curve) uninformative, in this study, we thus recorded F1 scores. In addition, we also performed bootstrapping (detailed in the following section) on the DNN-based classifier to obtain statistical robustness.

Several models, including ridge regression classifier (RR, scikit-learn RidgeClassifier), random forest (RF, scikit-learn RandomForestClassifier), linear support vector machine (LSVM, scikit-learn LinearSVC), gradient boosting (GB, scikit-learn GradientBoostingClassifier) and a dense neural network (DNN, built in keras) were implemented to manifest the classification. RR, RF, LSVM, and GB were conducted using default hyperparameters, except for class_weight, which was set to “balanced”. Details of DNN architecture, hyperparameters, and training are provided in **Table S13**.

#### Evaluation of the DNN-based classifier using bootstrapping

As a final model evaluation, we performed bootstrapping to generate 100 separate train-test splits, re-trained the model on each one, recorded individual F1 scores, and plotted the distribution of recorded F1 scores (**Fig. S5**).

#### Statistics

All statistical tests (**Fig. 4F**, **Fig. 5E**, and **5F**) were performed with R with default options. Details are provided where appropriate in the main text.

#### Code availability

We documented the complete PORT-EK pipeline and analytical scripts in Github, which is available for public download at https://github.com/wis-janusz/PORT-EK. The rest of the analytical scripts are available for public download at https://github.com/HCAngelC/K_mers_for_coronaviruses_cross_species_transmission.

#### Data availability

A list of metagenomic datasets used in this study is provided in **Data and materials availability**. Readouts from PORT-EK, including a list of enriched *k*-mers identified in the deer and bat datasets, coordinates of mutated amino acids coupled with nucleotide mutations present in enriched *k*-mers identified in the deer and bat datasets, recorded running times and peak memory usage for the datasets used in this study, and details of dense neural network (DNN) architecture, hyperparameters, and training is provided in **Tables S8-S13**, respectively and available for public download.

## Acknowledgments

We gratefully acknowledge all data contributors, the authors and their originating laboratories responsible for obtaining the specimens, and their submitting laboratories, for generating the genetic sequence and metadata and sharing via the GISAID Initiative, on which this study is based. We would like to thank G.J. Filion (University of Toronto Scarborough) for his proofreading and critical feedback on the manuscript.

## Author contributions

Conceptualization: HCC

Methodology: JW and HCC

Software: JW and HCC

Formal analysis: JW and HCC

Investigation, JW and HCC

Resources: HCC

Data curation: JW and HCC

Writing of original draft manuscript: JW and HCC

Writing, manuscript review and editing: JW and HCC

Visualization: JW and HCC

Supervision, HCC

Project administration, HCC

Funding acquisition, HCC

## Data and materials availability

The sequences used in this study are available from either the GISAID or NCBI Virus database as follows:

1. Deer dataset: US white-tailed deer group - EPI_SET_240422va, https://doi.org/10.55876/gis8.240422va (**Table S1**).
2. Deer dataset: early 2021 human group (April 2021) - EPI_SET_240422rw, https://doi.org/10.55876/gis8.240422rw (**Table S2**).
3. Deer set: late 2021 human group (November 2021) - EPI_SET_240422qc, https://doi.org/10.55876/gis8.240422qc (**Table S3**).
4. OoB dataset: US white-tailed deer group - EPI_SET_240422oy, https://doi.org/10.55876/gis8.240422oy (**Table S4**).
5. OoB dataset: human group - EPI_SET_240422xu, https://doi.org/10.55876/gis8.240422xu (**Table S5**).
6. Bat dataset: bat group- NCBI Virus accession numbers (**Table S6**).
7. Bat dataset, human group - EPI_SET_240422qm, https://doi.org/10.55876/gis8.240422qm (**Table S7**).

Of note, the supplementary tables for GISAID data are provided in the pdf format and used as is without any modification. All data generated by PORT-EK are available in the main text or the supplementary materials.

## Supplementary Materials

Supplementary Materials are referred to the separated file named **advances supplementary materials PORT-EK**.

## References

1. K. E. Jones, N. G. Patel, M. A. Levy, A. Storeygard, D. Balk, J. L. Gittleman, P. Daszak, Global trends in emerging infectious diseases. Nature 451, 990–993 (2008).

2. C. Drosten, S. Günther, W. Preiser, S. van der Werf, H.-R. Brodt, S. Becker, H. Rabenau, M. Panning, L. Kolesnikova, R. A. M. Fouchier, A. Berger, A.-M. Burguière, J. Cinatl, M. Eickmann, N. Escriou, K. Grywna, S. Kramme, J.-C. Manuguerra, S. Müller, V. Rickerts, M. Stürmer, S. Vieth, H.-D. Klenk, A. D. M. E. Osterhaus, H. Schmitz, H. W. Doerr, Identification of a novel coronavirus in patients with severe acute respiratory syndrome. N. Engl. J. Med. 348, 1967–1976 (2003).

3. T. G. Ksiazek, D. Erdman, C. S. Goldsmith, S. R. Zaki, T. Peret, S. Emery, S. Tong, C. Urbani, J. A. Comer, W. Lim, P. E. Rollin, S. F. Dowell, A.-E. Ling, C. D. Humphrey, W.-J. Shieh, J. Guarner, C. D. Paddock, P. Rota, B. Fields, J. DeRisi, J.-Y. Yang, N. Cox, J. M. Hughes, J. W. LeDuc, W. J. Bellini, L. J. Anderson, SARS Working Group, A novel coronavirus associated with severe acute respiratory syndrome. N. Engl. J. Med. 348, 1953–1966 (2003).

4. N. S. Zhong, B. J. Zheng, Y. M. Li, Poon, Z. H. Xie, K. H. Chan, P. H. Li, S. Y. Tan, Q. Chang, J. P. Xie, X. Q. Liu, J. Xu, D. X. Li, K. Y. Yuen, Peiris, Y. Guan, Epidemiology and cause of severe acute respiratory syndrome (SARS) in Guangdong, People’s Republic of China, in February, 2003. Lancet 362, 1353–1358 (2003).

5. S. Jiang, Z. Shi, Y. Shu, J. Song, G. F. Gao, W. Tan, D. Guo, A distinct name is needed for the new coronavirus. Lancet 395, 949 (2020).

6. N. Zhu, D. Zhang, W. Wang, X. Li, B. Yang, J. Song, X. Zhao, B. Huang, W. Shi, R. Lu, P. Niu, F. Zhan, X. Ma, D. Wang, W. Xu, G. Wu, G. F. Gao, W. Tan, China Novel Coronavirus Investigating and Research Team, A Novel Coronavirus from Patients with Pneumonia in China, 2019. N. Engl. J. Med. 382, 727–733 (2020).

7. M. Vignuzzi, J. K. Stone, J. J. Arnold, C. E. Cameron, R. Andino, Quasispecies diversity determines pathogenesis through cooperative interactions in a viral population. Nature 439, 344–348 (2006).

8. K. M. Peck, A. S. Lauring, Complexities of Viral Mutation Rates. J. Virol. 92 (2018).

9. J. E. Jones, V. Le Sage, S. S. Lakdawala, Viral and host heterogeneity and their effects on the viral life cycle. Nat. Rev. Microbiol. 19, 272–282 (2021).

10. T. P. Peacock, R. Penrice-Randal, J. A. Hiscox, W. S. Barclay, SARS-CoV-2 one year on: evidence for ongoing viral adaptation. J. Gen. Virol. 102 (2021).

11. K. M. Pepin, S. Lass, J. R. C. Pulliam, A. F. Read, J. O. Lloyd-Smith, Identifying genetic markers of adaptation for surveillance of viral host jumps. Nat. Rev. Microbiol. 8, 802–813 (2010).

12. J. M. Kirk, S. O. Kim, K. Inoue, M. J. Smola, D. M. Lee, M. D. Schertzer, J. S. Wooten, A. R. Baker, D. Sprague, D. W. Collins, C. R. Horning, S. Wang, Q. Chen, K. M. Weeks, P. J. Mucha, J. M. Calabrese, Functional classification of long non-coding RNAs by k-mer content. Nat. Genet. 50, 1474–1482 (2018).

13. C. Lorenzi, S. Barriere, J.-P. Villemin, L. Dejardin Bretones, A. Mancheron, W. Ritchie, iMOKA: k-mer based software to analyze large collections of sequencing data. Genome Biol. 21, 261 (2020).

14. D. R. Forsdyke, Success of alignment-free oligonucleotide (k-mer) analysis confirms relative importance of genomes not genes in speciation and phylogeny. Biol. J. Linn. Soc. Lond., doi: 10.1093/biolinnean/blz096 (2019).

15. W. Li, J. Freudenberg, J. Freudenberg, Alignment-free approaches for predicting novel Nuclear Mitochondrial Segments (NUMTs) in the human genome. Gene 691, 141–152 (2019).

16. E. Petrucci, L. Noé, C. Pizzi, M. Comin, “Iterative spaced seed hashing: Closing the gap between spaced seed hashing and k-mer hashing” in Bioinformatics Research and Applications (Springer International Publishing, Cham, 2019)Lecture notes in computer science, pp. 208–219.

17. Y. Ma, Z. Yu, R. Tang, X. Xie, G. Han, V. V. Anh, Phylogenetic Analysis of HIV-1 Genomes Based on the Position-Weighted K-mers Method. Entropy 22 (2020).

18. J. Wen, Y. Zhang, S. S. T. Yau, k-mer sparse matrix model for genetic sequence and its applications in sequence comparison. J. Theor. Biol. 363, 145–150 (2014).

19. A. B. Nassif, M. A. Talib, Q. Nasir, F. M. Dakalbab, Machine learning for anomaly detection: A systematic review. IEEE Access 9, 78658–78700 (2021).

20. H. Ren, Y. Li, T. Huang, Anomaly Detection Models for SARS-CoV-2 Surveillance Based on Genome -mers. Microorganisms 11 (2023).

21. K. J. V. Nordström, M. C. Albani, G. V. James, C. Gutjahr, B. Hartwig, F. Turck, U. Paszkowski, G. Coupland, K. Schneeberger, Mutation identification by direct comparison of whole-genome sequencing data from mutant and wild-type individuals using k-mers. Nat. Biotechnol. 31, 325–330 (2013).

22. N. L. Bray, H. Pimentel, P. Melsted, L. Pachter, Near-optimal probabilistic RNA-seq quantification. Nat. Biotechnol. 34, 525–527 (2016).

23. B. D. Ondov, T. J. Treangen, P. Melsted, A. B. Mallonee, N. H. Bergman, S. Koren, A. M. Phillippy, Mash: fast genome and metagenome distance estimation using MinHash. Genome Biol. 17, 132 (2016).

24. A. Shajii, D. Yorukoglu, Y. William Yu, B. Berger, Fast genotyping of known SNPs through approximate k-mer matching. Bioinformatics 32, i538–i544 (2016).

25. R. Patro, G. Duggal, M. I. Love, R. A. Irizarry, C. Kingsford, Salmon provides fast and bias-aware quantification of transcript expression. Nat. Methods 14, 417–419 (2017).

26. R. Ounit, S. Wanamaker, T. J. Close, S. Lonardi, CLARK: fast and accurate classification of metagenomic and genomic sequences using discriminative k-mers. BMC Genomics 16, 236 (2015).

27. F. P. Breitwieser, D. N. Baker, S. L. Salzberg, KrakenUniq: confident and fast metagenomics classification using unique k-mer counts. Genome Biol. 19, 198 (2018).

28. J. Audoux, N. Philippe, R. Chikhi, M. Salson, M. Gallopin, M. Gabriel, J. Le Coz, E. Drouineau, T. Commes, D. Gautheret, DE-kupl: exhaustive capture of biological variation in RNA-seq data through k-mer decomposition. Genome Biol. 18, 243 (2017).

29. B. T. Lau, D. Pavlichin, A. C. Hooker, A. Almeda, G. Shin, J. Chen, M. K. Sahoo, C. H. Huang, B. A. Pinsky, H. J. Lee, H. P. Ji, Profiling SARS-CoV-2 mutation fingerprints that range from the viral pangenome to individual infection quasispecies. Genome Med. 13, 62 (2021).

30. S. C. Manekar, S. R. Sathe, A benchmark study of k-mer counting methods for high-throughput sequencing. Gigascience 7 (2018).

31. P. Creixell, E. M. Schoof, C. S. H. Tan, R. Linding, Mutational properties of amino acid residues: implications for evolvability of phosphorylatable residues. Philos. Trans. R. Soc. Lond. B Biol. Sci. 367, 2584–2593 (2012).

32. G. Louppe, Understanding random forests: From theory to practice. doi: 10.48550/ARXIV.1407.7502 (2014).

33. X. Li, Y. Wang, S. Basu, K. Kumbier, B. Yu, A debiased MDI feature importance measure for Random Forests. doi: 10.48550/ARXIV.1906.10845 (2019).

34. J. H. Moore, Bootstrapping, permutation testing and the method of surrogate data. Phys. Med. Biol. 44, L11–2 (1999).

35. R. Gibb, D. W. Redding, K. Q. Chin, C. A. Donnelly, T. M. Blackburn, T. Newbold, K. E. Jones, Zoonotic host diversity increases in human-dominated ecosystems. Nature 584, 398–402 (2020).

36. N. J. Loman, R. V. Misra, T. J. Dallman, C. Constantinidou, S. E. Gharbia, J. Wain, M. J. Pallen, Performance comparison of benchtop high-throughput sequencing platforms. Nat. Biotechnol. 30, 434–439 (2012).

37. S. Jünemann, F. J. Sedlazeck, K. Prior, A. Albersmeier, U. John, J. Kalinowski, A. Mellmann, A. Goesmann, A. von Haeseler, J. Stoye, D. Harmsen, Updating benchtop sequencing performance comparison, Nature biotechnology. 31 (2013)pp. 294–296.

38. F. Meacham, D. Boffelli, J. Dhahbi, D. I. K. Martin, M. Singer, L. Pachter, Identification and correction of systematic error in high-throughput sequence data. BMC Bioinformatics 12, 451 (2011).

39. A. M. Carabelli, T. P. Peacock, L. G. Thorne, W. T. Harvey, J. Hughes, COVID-19 Genomics UK Consortium, S. J. Peacock, W. S. Barclay, T. I. de Silva, G. J. Towers, D. L. Robertson, SARS-CoV-2 variant biology: immune escape, transmission and fitness. Nat. Rev. Microbiol. 21, 162–177 (2023).

40. E. Volz, V. Hill, J. T. McCrone, A. Price, D. Jorgensen, Á. O’Toole, J. Southgate, R. Johnson, B. Jackson, F. F. Nascimento, S. M. Rey, S. M. Nicholls, R. M. Colquhoun, A. da Silva Filipe, J. Shepherd, D. J. Pascall, R. Shah, N. Jesudason, K. Li, R. Jarrett, N. Pacchiarini, M. Bull, L. Geidelberg, I. Siveroni, COG-UK Consortium, I. Goodfellow, N. J. Loman, O. G. Pybus, D. L. Robertson, E. C. Thomson, A. Rambaut, T. R. Connor, Evaluating the Effects of SARS-CoV-2 Spike Mutation D614G on Transmissibility and Pathogenicity. Cell 184, 64–75.e11 (2021).

41. Á. O’Toole, O. G. Pybus, M. E. Abram, E. J. Kelly, A. Rambaut, Pango lineage designation and assignment using SARS-CoV-2 spike gene nucleotide sequences. BMC Genomics 23, 121 (2022).

42. J. L. Chaney, P. L. Clark, Roles for Synonymous Codon Usage in Protein Biogenesis. Annu. Rev. Biophys. 44, 143–166 (2015).

43. P. M. Sharp, E. Cowe, D. G. Higgins, D. C. Shields, K. H. Wolfe, F. Wright, Codon usage patterns in Escherichia coli, Bacillus subtilis, Saccharomyces cerevisiae, Schizosaccharomyces pombe, Drosophila melanogaster and Homo sapiens; a review of the considerable within-species diversity. Nucleic Acids Res. 16, 8207–8211 (1988).

44. I. Bahir, M. Fromer, Y. Prat, M. Linial, Viral adaptation to host: a proteome-based analysis of codon usage and amino acid preferences. Mol. Syst. Biol. 5, 311 (2009).

45. M. Costantini, R. Cammarano, G. Bernardi, The evolution of isochore patterns in vertebrate genomes. BMC Genomics 10, 146 (2009).

46. G. Bernardi, G. Bernardi, Compositional constraints and genome evolution. J. Mol. Evol. 24, 1–11 (1986).

47. L. Duret, Evolution of synonymous codon usage in metazoans. Curr. Opin. Genet. Dev. 12, 640–649 (2002).

48. A. Carbone, Codon bias is a major factor explaining phage evolution in translationally biased hosts. J. Mol. Evol. 66, 210–223 (2008).

49. J. W. Barrett, Y. Sun, S. H. Nazarian, T. A. Belsito, C. R. Brunetti, G. McFadden, Optimization of codon usage of poxvirus genes allows for improved transient expression in mammalian cells. Virus Genes 33, 15– 26 (2006).

50. L. A. Shackelton, C. R. Parrish, E. C. Holmes, Evolutionary basis of codon usage and nucleotide composition bias in vertebrate DNA viruses. J. Mol. Evol. 62, 551–563 (2006).

51. K.-N. Zhao, W. J. Liu, I. H. Frazer, Codon usage bias and A+T content variation in human papillomavirus genomes. Virus Res. 98, 95–104 (2003).

52. B. Berkhout, A. Grigoriev, M. Bakker, V. V. Lukashov, Codon and amino acid usage in retroviral genomes is consistent with virus-specific nucleotide pressure. AIDS Res. Hum. Retroviruses 18, 133–141 (2002).

53. A. M. Alonso, L. Diambra, SARS-CoV-2 Codon Usage Bias Downregulates Host Expressed Genes With Similar Codon Usage. Front Cell Dev Biol 8, 831 (2020).

54. W. Hou, Characterization of codon usage pattern in SARS-CoV-2. Virol. J. 17, 138 (2020).

55. W. Gu, T. Zhou, J. Ma, X. Sun, Z. Lu, Analysis of synonymous codon usage in SARS Coronavirus and other viruses in the Nidovirales. Virus Res. 101, 155–161 (2004).

56. V. Presnyak, N. Alhusaini, Y.-H. Chen, S. Martin, N. Morris, N. Kline, S. Olson, D. Weinberg, K. E. Baker, B. R. Graveley, J. Coller, Codon optimality is a major determinant of mRNA stability. Cell 160, 1111– 1124 (2015).

57. R. L. Carneiro, R. D. Requião, S. Rossetto, T. Domitrovic, F. L. Palhano, Codon stabilization coefficient as a metric to gain insights into mRNA stability and codon bias and their relationships with translation. Nucleic Acids Res. 47, 2216–2228 (2019).

58. C. J. Epstein, Non-randomness of amino-acid changes in the evolution of homologous proteins. Nature 215, 355–359 (1967).

59. N. Goldman, Z. Yang, A codon-based model of nucleotide substitution for protein-coding DNA sequences. Mol. Biol. Evol. 11, 725–736 (1994).

60. J. Zhang, Rates of conservative and radical nonsynonymous nucleotide substitutions in mammalian nuclear genes. J. Mol. Evol. 50, 56–68 (2000).

61. N. G. C. Smith, Are radical and conservative substitution rates useful statistics in molecular evolution? J. Mol. Evol. 57, 467–478 (2003).

62. K. Popadin, L. V. Polishchuk, L. Mamirova, D. Knorre, K. Gunbin, Accumulation of slightly deleterious mutations in mitochondrial protein-coding genes of large versus small mammals. Proc. Natl. Acad. Sci. U. S. A. 104, 13390–13395 (2007).

63. C. C. Weber, B. Nabholz, J. Romiguier, H. Ellegren, Kr/Kc but not dN/dS correlates positively with body mass in birds, raising implications for inferring lineage-specific selection. Genome Biol. 15, 542 (2014).

64. C. C. Weber, S. Whelan, Physicochemical Amino Acid Properties Better Describe Substitution Rates in Large Populations. Mol. Biol. Evol. 36, 679–690 (2019).

65. E. Domingo, J. Sheldon, C. Perales, Viral quasispecies evolution. Microbiol. Mol. Biol. Rev. 76, 159–216 (2012).

66. E. Domingo, C. Perales, Viral quasispecies. PLoS Genet. 15, e1008271 (2019).

67. S. Chen, C. He, Y. Li, Z. Li, C. E. Melançon, A computational toolset for rapid identification of SARS-CoV-2, other viruses and microorganisms from sequencing data. Brief. Bioinform. 22, 924–935 (2021).

68. R. Islam, R. S. Raju, N. Tasnim, I. H. Shihab, M. A. Bhuiyan, Y. Araf, T. Islam, Choice of assemblers has a critical impact on de novo assembly of SARS-CoV-2 genome and characterizing variants. Brief. Bioinform. 22 (2021).

69. S. Pei, S. S.-T. Yau, Analysis of the Genomic Distance Between Bat Coronavirus RaTG13 and SARS-CoV-2 Reveals Multiple Origins of COVID-19. Acta Math Sci 41, 1017–1022 (2021).

70. J. Avila Cartes, S. Anand, S. Ciccolella, P. Bonizzoni, G. Della Vedova, Accurate and fast clade assignment via deep learning and frequency chaos game representation. Gigascience 12 (2022).

71. G. Nicora, M. Salemi, S. Marini, R. Bellazzi, Predicting emerging SARS-CoV-2 variants of concern through a One Class dynamic anomaly detection algorithm. BMJ Health Care Inform 29 (2022).

72. R. S. Raju, A. Al Nahid, P. Chondrow Dev, R. Islam, VirusTaxo: Taxonomic classification of viruses from the genome sequence using k-mer enrichment. Genomics 114, 110414 (2022).

73. I. Sung, S. Lee, M. Pak, Y. Shin, S. Kim, AutoCoV: tracking the early spread of COVID-19 in terms of the spatial and temporal patterns from embedding space by K-mer based deep learning. BMC Bioinformatics 23, 149 (2022).

74. K. Walker, D. Kalra, R. Lowdon, G. Chen, D. Molik, D. C. Soto, F. Dabbaghie, A. A. Khleifat, M. Mahmoud, L. F. Paulin, M. S. Raza, S. P. Pfeifer, D. P. Agustinho, E. Aliyev, P. Avdeyev, E. R. Barrozo, S. Behera, K. Billingsley, L. C. Chong, D. Choubey, W. De Coster, Y. Fu, A. R. Gener, T. Hefferon, D. M. Henke, W. Höps, A. Illarionova, M. D. Jochum, M. Jose, R. K. Kesharwani, S. R. R. Kolora, J. Kubica, P. Lakra, D. Lattimer, C.-S. Liew, B.-W. Lo, C. Lo, A. Lötter, S. Majidian, S. K. Mendem, R. Mondal, H. Ohmiya, N. Parvin, C. Peralta, C.-L. Poon, R. Prabhakaran, M. Saitou, A. Sammi, P. Sanio, N. Sapoval, N. Syed, T. Treangen, G. Wang, T. Xu, J. Yang, S. Zhang, W. Zhou, F. J. Sedlazeck, B. Busby, The third international hackathon for applying insights into large-scale genomic composition to use cases in a wide range of organisms. F1000Res. 11, 530 (2022).

75. R. Chandra, C. Bansal, M. Kang, T. Blau, V. Agarwal, P. Singh, L. O. W. Wilson, S. Vasan, Unsupervised machine learning framework for discriminating major variants of concern during COVID-19. PLoS One 18, e0285719 (2023).

76. A. S. Thind, S. Sinha, Using Chaos-Game-Representation for Analysing the SARS-CoV-2 Lineages, Newly Emerging Strains and Recombinants. Curr. Genomics 24, 187–195 (2023).

77. A. Thommana, M. Shakya, J. Gandhi, C. K. Fung, P. S. G. Chain, I. Maljkovic Berry, M. A. Conte, Intrahost SARS-CoV-2 k-mer Identification Method (iSKIM) for Rapid Detection of Mutations of Concern Reveals Emergence of Global Mutation Patterns. Viruses 14 (2022).

78. Y. W. Yu, On Minimizers and Convolutional Filters: Theoretical Connections and Applications to Genome Analysis. J. Comput. Biol. 31, 381–395 (2024).

79. S. Khare, C. Gurry, L. Freitas, M. B. Schultz, G. Bach, A. Diallo, N. Akite, J. Ho, R. T. Lee, W. Yeo, G. C. Curation Team, S. Maurer-Stroh, GISAID’s Role in Pandemic Response. China CDC Wkly 3, 1049–1051 (2021).

80. E. L. Hatcher, S. A. Zhdanov, Y. Bao, O. Blinkova, E. P. Nawrocki, Y. Ostapchuck, A. A. Schäffer, J. R. Brister, Virus Variation Resource - improved response to emergent viral outbreaks. Nucleic Acids Res. 45, D482–D490 (2017).

81. Z. Gu, R. Eils, M. Schlesner, Complex heatmaps reveal patterns and correlations in multidimensional genomic data. Bioinformatics 32, 2847–2849 (2016).

82. Z. Gu, Complex heatmap visualization. Imeta 1 (2022).

83. The pandas development team, Pandas-Dev/pandas: Pandas (Zenodo, 2024; https://zenodo.org/doi/10.5281/zenodo.3509134).

84. A. Altmann, L. Toloşi, O. Sander, T. Lengauer, Permutation importance: a corrected feature importance measure. Bioinformatics 26, 1340–1347 (2010).

85. Y. Wang, Q. Chen, C. Deng, Y. Zheng, F. Sun, KmerGO: A Tool to Identify Group-Specific Sequences With -mers. Front. Microbiol. 11, 2067 (2020).

