## Supplementary Figures and Tables for "Identification of potential SARS-CoV-2 genetic markers resulting from host domestication"

**This PDF file includes:**

Supplementary Text
Figs. S1 to S6
Tables S1 to S13

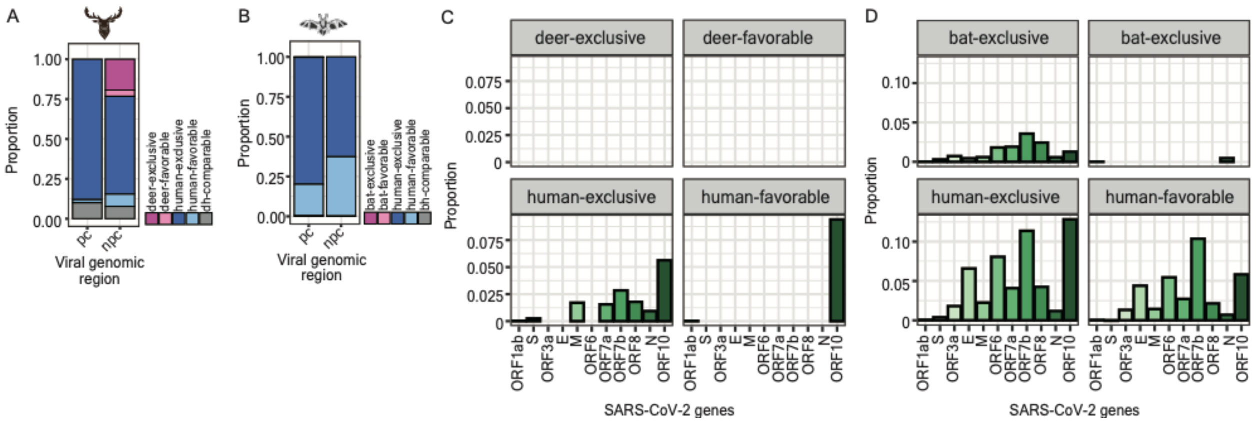

**Fig. S1.**

**Landscapes of enriched *k*-mers across the SARS-CoV-2 genome.** (**A**, **B**) Stacked bar charts representing the proportion of enriched *k*-mers that overlay with the protein-coding (pc) and non-protein coding (npc) regions throughout the SARS-CoV-2 genome (**A**, enriched *k*-mers identified in the deer dataset; **B**, enriched *k*-mers identified in the bat dataset). The dark pink color indicates the SARS-CoV-2 genomic loci overlaid exclusively with enriched *k*-mers identified in deer (**A**) or bat (**B**) isolates; The light pink color indicates the loci overlaid preferentially with enriched *k*-mers identified in deer (**A**) or bat (**B**) isolates; the dark blue color indicates the SARS-CoV-2 genomic loci overlaid exclusively with enriched *k*-mers identified in human isolates collected in the deer (**A**) or bat (**B**) dataset; the light blue color indicates the SARS-CoV-2 genomic loci overlaid preferentially with enriched *k*-mers identified in human isolates collected in the deer (**A**) or bat (**B**) dataset; the grey color indicates the loci targeted by the identical number of enriched *k*-mers between animal (deer and bat) and human isolates. (**C**, **D**) Bar charts representing the enrichment of enriched *k*-mers identified in the deer (**C**) and bat (**D**) datasets per SARS-CoV-2 gene. The calculation of the enrichment is described in the **Materials and Methods**. Four facets in the panel (**C**) and (**D**) are separated based on the intrinsic property of a genomic locus designated as deer- (**C**) or bat- (**D**) exclusive sites or deer- (**C**) or bat- (**D**) favorable sites. Details are referred to in the main text.

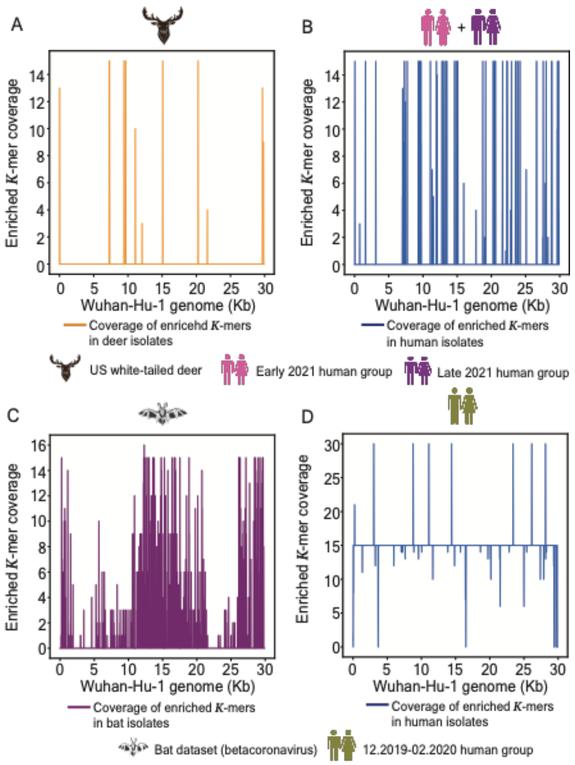

**Fig. S2.**

**The coverage landscapes of enriched *k*-mers across the SARS-CoV-2 genome.** (**A**, **B**, **C**, **D**) Line plots representing the coverage of the SARS-CoV-2 genome per site overlaid with enriched *k*-mers identified in deer (**A**) and human (**B**) isolates collected in the deer dataset or bat (**C**) and human (**D**) isolates collected in the bat dataset.

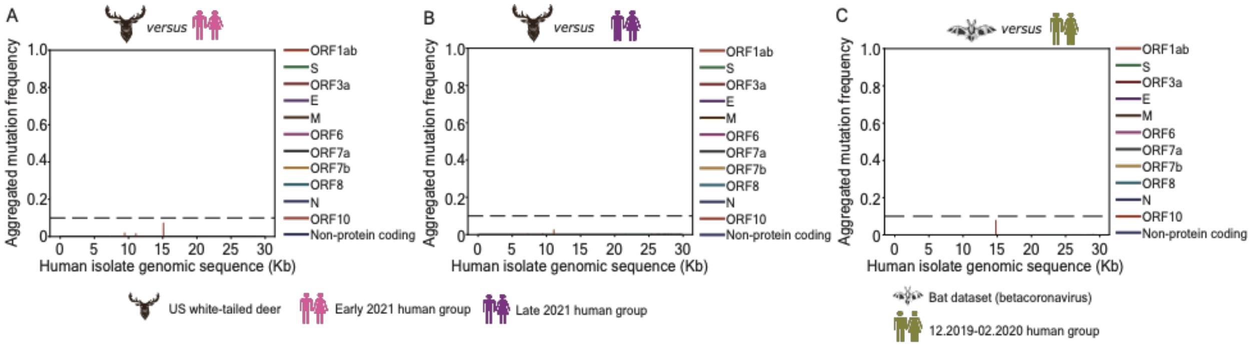

**Fig. S3.**

**Verification of the specificity of mutations emerging from the enriched *k*-mers identified in animal isolates.** (**A**, **B**) Line plots representing the frequency of the presence of mutations detected from the enriched *k*-mers in deer isolates against the genomic sequences of human isolates collected in the early (**A**) and late (**B**) 2021 human groups. (**C**) Line plots representing the frequency of the presence of mutations detected from the enriched *k*-mers in bat isolates against the genomic sequences of human isolates collected in the bat dataset.

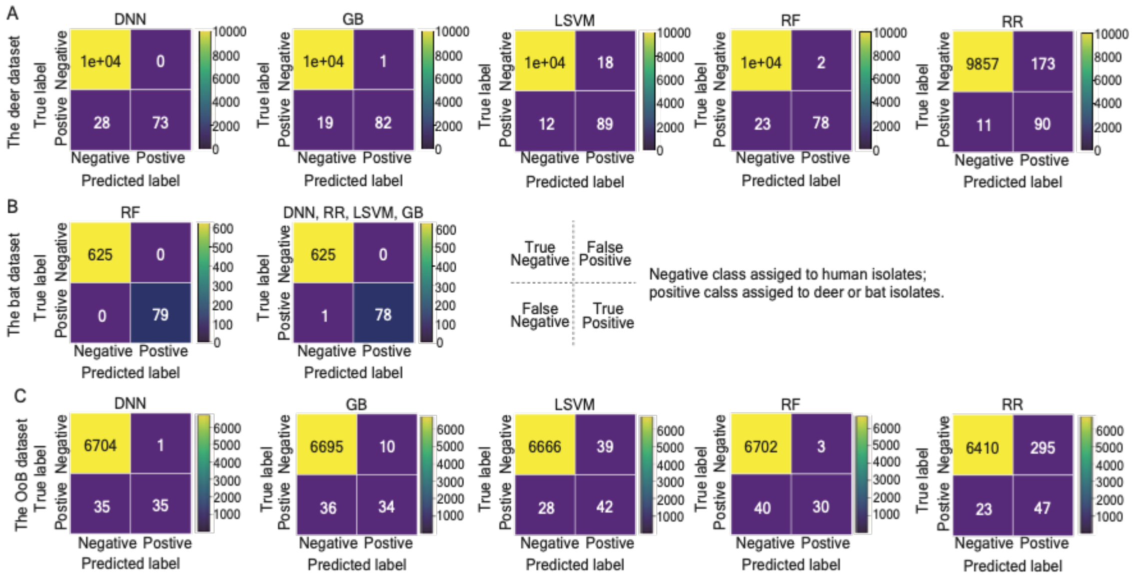

**Fig. S4.**

**Prediction summary of the robustness of testing classifiers subjected to different model architectures.** Confusion matrices summarizing the number of predictions that are true negative (top corner on the left-hand side), true positive (bottom corner on the right-hand side), false negative (bottom corner on the left-hand side), and false positive (top corner on the right-hand side). The negative class is designated to individual SARS-CoV-2 genomes predicted as human isolates; the positive class is designated to individual SARS-CoV-2 genomes predicted as animal (deer or bats) isolates. Classifiers were constructed based on the total count of enriched *k*-mers and subjected to the following models, dense neural network (DNN), gradient boosting (GB), linear support vector machine (LSVM), random forest (RF), ridge regression (RR). We applied them to predict the most probable host species of individual SARS-CoV-2 genomic sequences collected in the deer (**A**), bat (**B**), and out of the bag (OoB) datasets (**C**).

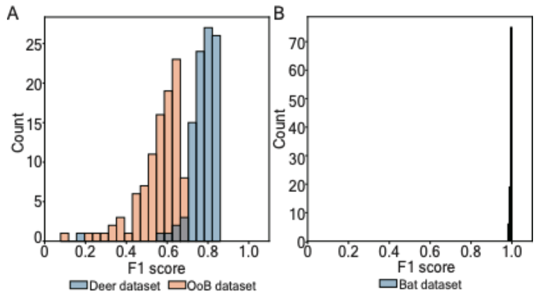

**Fig. S5.**

**Bootstrapping on the subsets of metagenomes in testing datasets.** (**A**, **B**) Histogram representing the distribution of F1 scores measured from 100 independent train-test splits on single genomes of isolates collected in the deer, OoB (**A**), and bat (**B**) datasets.

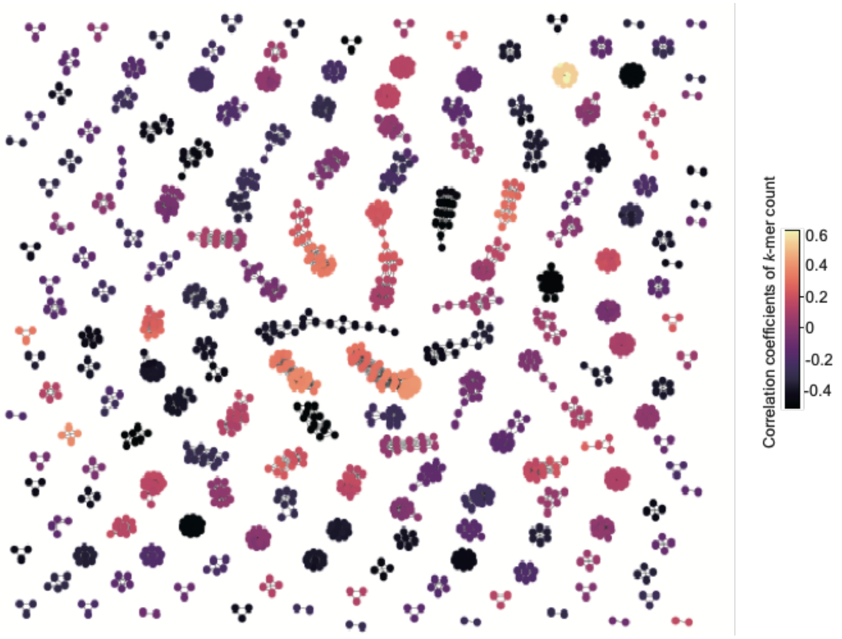

**Fig. S6.**

**Correlation distribution of a subset of enriched *k*-mers identified in the deer dataset based on their counts.** Graph network representing a correlation of the groups of *k*-mers with a high correlation of their counts. Nodes represent individual enriched *k*-mers; edges represent a correlation coefficient of more than 0.9 between two adjacent nodes. The color scale represents the absolute value of the correlation coefficient computed based on the count of enriched *k*-mers.

**Table S1.**

Deer dataset: US white-tailed deer group - EPI_SET_240422va, https://doi.org/10.55876/gis8.240422va.

GISIAD supplemental table for deer coronavirus sequences of deer dataset.

SUPPLEMENTAL TABLE

**Data Availability**

GISAID Identifier: EPI_SET_240422va
doi: 10.55876/gis8.240422va

All genome sequences and associated metadata in this dataset are published in GISAID's EpiCoV database. To view the contributors of each individual sequence with details such as accession number, Virus name, Collection date, Originating Lab and Submitting Lab and the list of Authors, visit 10.55876/gis8.240422va

**Data Snapshot**

- EPI_SET_240422va is composed of 336 individual genome sequences.
- The collection dates range from 2021-10-01 to 2021-12-30;
- Data were collected in 2 countries and territories;
- All sequences in this dataset are compared relative to hCoV-19/Wuhan/WIV04/2019 (WIV04), the official reference sequence employed by GISAID (EPI_ISL_402124). Learn more at https://gisaid.org/WIV04.

**Table S2.**

Deer dataset: early 2021 human group (April 2021) - EPI_SET_240422rw, https://doi.org/10.55876/gis8.240422rw.

GISIAD supplemental table for early 2021 human coronavirus sequences of deer dataset.

SUPPLEMENTAL TABLE

**Data Availability**

GISAID Identifier: EPI_SET_240422rw
doi: 10.55876/gis8.240422rw

All genome sequences and associated metadata in this dataset are published in GISAID's EpiCoV database. To view the contributors of each individual sequence with details such as accession number, Virus name, Collection date, Originating Lab and Submitting Lab and the list of Authors, visit 10.55876/gis8.240422rw

**Data Snapshot**

- EPI_SET_240422rw is composed of 21,905 individual genome sequences.
- The collection dates range from 2021-04-05 to 2021-04-09;
- Data were collected in 1 countries and territories;
- All sequences in this dataset are compared relative to hCoV-19/Wuhan/WIV04/2019 (WIV04), the official reference sequence employed by GISAID (EPI_ISL_402124). Learn more at https://gisaid.org/WIV04.

**Table S3.**

Deer set: late 2021 human group (November 2021) - EPI_SET_240422qc, https://doi.org/10.55876/gis8.240422qc.

GISIAD supplemental table for late 2021 human coronavirus sequences of deer dataset.

SUPPLEMENTAL TABLE

**Data Availability**

GISAID Identifier: EPI_SET_240422qc
doi: 10.55876/gis8.240422qc

All genome sequences and associated metadata in this dataset are published in GISAID's EpiCoV database. To view the contributors of each individual sequence with details such as accession number, Virus name, Collection date, Originating Lab and Submitting Lab and the list of Authors, visit 10.55876/gis8.240422qc

**Data Snapshot**

- EPI_SET_240422qc is composed of 11,526 individual genome sequences.
- The collection dates range from 2021-11-14 to 2021-11-15;
- Data were collected in 1 countries and territories;
- All sequences in this dataset are compared relative to hCoV-19/Wuhan/WIV04/2019 (WIV04), the official reference sequence employed by GISAID (EPI_ISL_402124). Learn more at https://gisaid.org/WIV04.

**Table S4.**

OoB dataset: US white-tailed deer group - EPI_SET_240422oy, https://doi.org/10.55876/gis8.240422oy.

GISIAD supplemental table for deer coronavirus sequences of OoB dataset.

SUPPLEMENTAL TABLE

**Data Availability**

GISAID Identifier: EPI_SET_240422oy
doi: 10.55876/gis8.240422oy

All genome sequences and associated metadata in this dataset are published in GISAID's EpiCoV database. To view the contributors of each individual sequence with details such as accession number, Virus name, Collection date, Originating Lab and Submitting Lab and the list of Authors, visit 10.55876/gis8.240422oy

**Data Snapshot**

- EPI_SET_240422oy is composed of 70 individual genome sequences.
- The collection dates range from 2022-01-03 to 2023-11-10;
- Data were collected in 1 countries and territories;
- All sequences in this dataset are compared relative to hCoV-19/Wuhan/WIV04/2019 (WIV04), the official reference sequence employed by GISAID (EPI_ISL_402124). Learn more at https://gisaid.org/WIV04.

**Table S5.**

OoB dataset: human group - EPI_SET_240422xu, https://doi.org/10.55876/gis8.240422xu.
GISIAD supplemental table for human coronavirus sequences of OoB dataset.

SUPPLEMENTAL TABLE

**Data Availability**

GISAID Identifier: EPI_SET_240422xu
doi: 10.55876/gis8.240422xu

All genome sequences and associated metadata in this dataset are published in GISAID's EpiCoV database. To view the contributors of each individual sequence with details such as accession number, Virus name, Collection date, Originating Lab and Submitting Lab and the list of Authors, visit 10.55876/gis8.240422xu

**Data Snapshot**

- EPI_SET_240422xu is composed of 6,705 individual genome sequences.
- The collection dates range from 2022-01-01 to 2022-01-15;
- Data were collected in 3 countries and territories;
- All sequences in this dataset are compared relative to hCoV-19/Wuhan/WIV04/2019 (WIV04), the official reference sequence employed by GISAID (EPI_ISL_402124). Learn more at https://gisaid.org/WIV04.

**Table S6.**

Bat dataset: bat group - NCBI Virus accession numbers.

NCBI Virus accession numbers of bat coronavirus sequences used in the bat dataset.

| |
|---|
| NC_030886.1 |
| NC_025217.1 |
| NC_014470.1 |
| NC_009019.1 |
| NC_009020.1 |
| NC_009021.1 |
| OR261262.1 |
| OR261263.1 |
| OR261264.1 |
| OR261265.1 |
| OR261266.1 |
| OR261267.1 |
| OR261268.1 |
| OR233291.1 |
| OR233292.1 |
| OR233293.1 |
| OR233294.1 |
| OR233295.1 |
| OR233296.1 |
| OR233297.1 |
| OR233298.1 |
| OR233299.1 |
| OR233300.1 |
| OR233301.1 |
| OR233302.1 |

| |
|---|
| OR233303.1 |
| OR233304.1 |
| OR233305.1 |
| OR233308.1 |
| OR233310.1 |
| OR233311.1 |
| OR233312.1 |
| OR233313.1 |
| OR233314.1 |
| OR233316.1 |
| OR233317.1 |
| OR233318.1 |
| OR233319.1 |
| OR233320.1 |
| OR233321.1 |
| OR233322.1 |
| OR233323.1 |
| OR233324.1 |
| OR233325.1 |
| OR233328.1 |
| OQ503495.1 |
| OQ503496.1 |
| OQ503497.1 |
| OQ503498.1 |
| OQ503499.1 |
| OQ503500.1 |
| OQ503501.1 |

| |
|---|
| OQ503502.1 |
| OQ503503.1 |
| OQ503504.1 |
| OQ503505.1 |
| OQ503506.1 |
| OK017908.1 |
| OQ401247.1 |
| OQ401248.1 |
| OQ401249.1 |
| OQ401251.1 |
| OQ405399.1 |
| OP776338.1 |
| OP776339.1 |
| OP776340.1 |
| OM995890.1 |
| OM995891.1 |
| LC663783.1 |
| LC706863.1 |
| LC706864.1 |
| LC706865.1 |
| LC663793.1 |
| OK287354.1 |
| OK287355.1 |
| ON325306.1 |
| ON325307.1 |
| MZ293739.1 |
| MZ293752.1 |

| |
|---|
| MZ293753.1 |
| MZ293754.1 |
| MZ293755.1 |
| MZ293756.1 |
| ON378802.1 |
| ON378807.1 |
| OM240725.1 |
| MW681002.1 |
| OK067319.1 |
| OK067320.1 |
| OK067321.1 |
| OL674074.1 |
| OL674075.1 |
| OL674076.1 |
| OL674077.1 |
| OL674078.1 |
| OL674079.1 |
| OL674080.1 |
| OL674081.1 |
| MZ328294.1 |
| MZ328297.1 |
| OK017792.1 |
| OK017793.1 |
| OK017794.1 |
| OK017795.1 |
| OK017796.1 |
| OK017797.1 |

| |
|---|
| OK017798.1 |
| OK017799.1 |
| OK017800.1 |
| OK017801.1 |
| OK017802.1 |
| OK017803.1 |
| OK017804.1 |
| OK017805.1 |
| OK017806.1 |
| OK017807.1 |
| OK017808.1 |
| OK017809.1 |
| OK017810.1 |
| OK017811.1 |
| OK017812.1 |
| OK017813.1 |
| OK017814.1 |
| OK017815.1 |
| OK017816.1 |
| OK017817.1 |
| OK017818.1 |
| OK017819.1 |
| OK017820.1 |
| OK017821.1 |
| OK017822.1 |
| OK017823.1 |
| OK017824.1 |

| |
|---|
| OK017825.1 |
| OK017826.1 |
| OK017827.1 |
| OK017828.1 |
| OK017829.1 |
| OK017830.1 |
| OK017831.1 |
| OK017832.1 |
| OK017833.1 |
| OK017834.1 |
| OK017835.1 |
| OK017836.1 |
| OK017837.1 |
| OK017838.1 |
| OK017839.1 |
| OK017840.1 |
| OK017841.1 |
| OK017842.1 |
| OK017843.1 |
| OK017844.1 |
| OK017845.1 |
| OK017846.1 |
| OK017847.1 |
| OK017848.1 |
| OK017849.1 |
| OK017850.1 |
| OK017851.1 |

| |
|---|
| OK017852.1 |
| OK017853.1 |
| OK017854.1 |
| OK017855.1 |
| OK017856.1 |
| OK017857.1 |
| OK017858.1 |
| OK017859.1 |
| OK017860.1 |
| MW719567.1 |
| MZ081376.1 |
| MZ081377.1 |
| MZ081378.1 |
| MZ081379.1 |
| MZ081380.1 |
| MZ081381.1 |
| MZ081382.1 |
| MZ190137.1 |
| MZ190138.1 |
| MT726043.1 |
| MT726045.1 |
| MW703458.1 |
| MW251308.1 |
| MW218395.1 |
| LC556375.1 |
| MT350598.1 |
| MN996532.2 |

| |
|---|
| MN611519.1 |
| MN611520.1 |
| MK211374.1 |
| MK211375.1 |
| MK211376.1 |
| MK211377.1 |
| MK211378.1 |
| MK211379.1 |
| KY352407.1 |
| MF593268.1 |
| MG772933.1 |
| MG772934.1 |
| MG762674.1 |
| KX442564.1 |
| KX442565.1 |
| MG596802.1 |
| MG596803.1 |
| KY417142.1 |
| KY417143.1 |
| KY417144.1 |
| KY417145.1 |
| KY417146.1 |
| KY417147.1 |
| KY417148.1 |
| KY417149.1 |
| KY417150.1 |
| KY417151.1 |

| |
|---|
| KY417152.1 |
| KY938558.1 |
| KU973692.1 |
| KY770858.1 |
| KY770859.1 |
| KY770860.1 |
| KU762338.1 |
| KU182964.1 |
| KT444582.1 |
| KJ473811.1 |
| KJ473812.1 |
| KJ473813.1 |
| KJ473814.1 |
| KJ473815.1 |
| KJ473816.1 |
| KJ473820.1 |
| KJ473822.1 |
| KP886808.1 |
| KP886809.1 |
| KF636752.1 |
| KJ473821.1 |
| KF569996.1 |
| KC881005.1 |
| KC881006.1 |
| KF367457.1 |
| KC869678.4 |
| JX993987.1 |

| |
|---|
| JX993988.1 |
| HM211098.1 |
| HM211099.1 |
| HM211100.1 |
| HM211101.1 |
| GU190215.1 |
| FJ588686.1 |
| EF065505.1 |
| EF065506.1 |
| EF065507.1 |
| EF065508.1 |
| EF065509.1 |
| EF065510.1 |
| EF065511.1 |
| EF065512.1 |
| EF065513.1 |
| EF065514.1 |
| EF065515.1 |
| EF065516.1 |
| DQ412042.1 |
| DQ412043.1 |
| DQ071615.1 |

**Table S7.**

Bat dataset, human group - EPI_SET_240422qm, https://doi.org/10.55876/gis8.240422qm. GISIAD supplemental table for human coronavirus sequences of bat dataset.

SUPPLEMENTAL TABLE

**Data Availability**

GISAID Identifier: EPI_SET_240422qm
doi: 10.55876/gis8.240422qm

All genome sequences and associated metadata in this dataset are published in GISAID's EpiCoV database. To view the contributors of each individual sequence with details such as accession number, Virus name, Collection date, Originating Lab and Submitting Lab and the list of Authors, visit 10.55876/gis8.240422qm

**Data Snapshot**

- EPI_SET_240422qm is composed of 2,081 individual genome sequences.
- The collection dates range from 2019-12-24 to 2020-02-29;
- Data were collected in 54 countries and territories;
- All sequences in this dataset are compared relative to hCoV-19/Wuhan/WIV04/2019 (WIV04), the official reference sequence employed by GISAID (EPI_ISL_402124). Learn more at https://gisaid.org/WIV04.

**Table S8.**

A list of enriched *k*-mers identified in the deer dataset.
Table containing sequences and statistics of *k*-mers enriched in deer dataset.

Please note that **Table S8** is provided separately due to the large dataset involved.

**Table S9.**

A list of enriched *k*-mers identified in the bat dataset.
Table containing sequences and statistics of *k*-mers enriched in the bat dataset.

Please note that **Table S9** is provided separately due to the large dataset involved.

**Table S10.**

Coordinates of mutated amino acids coupled with nucleotide mutations present in enriched *k*-mers identified in the deer dataset.

| mutation | type | ref_nucleotide | ref_pos | mut_nucleotide | n_kmers | max_deer_RMSE | agg_mutation_freq | early_agg_mutation_freq | late_agg_mutation_freq | group | gene | ref_codon | ref_amino_acid | mut_codon | mut_amino_acid | Protein |
|---|---|---|---|---|---|---|---|---|---|---|---|---|---|---|---|---|
| C7267T | enriched | C | 7267 | T | 15 | 0,1564 | 0,1577 | 0,0007 | 0,0021 | eq_cov | orf1ab | CAC | His | TAC | Cys | nsp3 |
| C7303T | enriched | C | 7303 | T | 15 | 0,5633 | 0,5655 | 0,0029 | 0,0015 | eq_cov | orf1ab | CAT | His | TAT | Cys | nsp3 |
| C9430T | enriched | C | 9430 | T | 15 | 0,4917 | 0,5030 | 0,0189 | 0,0039 | eq_cov | orf1ab | CGT | Arg | TGT | Cys | nsp4 |
| C9679T | enriched | C | 9679 | T | 15 | 0,1546 | 0,1548 | 5E-05 | 0,0003 | eq_cov | orf1ab | CTG | Leu | TTG | Leu | nsp4 |
| G11083T | enriched | G | 11083 | T | 10 | 0,1265 | 0,1458 | 0,0145 | 0,0246 | dep_pref | orf1ab | GTA | Val | TTA | Leu | nsp6 |
| T15096C | enriched | T | 15096 | C | 15 | 0,1040 | 0,1339 | 0,0724 | 0,0003 | eq_cov | orf1ab | AAT | Asn | AAC | Asn | RDRP |
| C20259T | enriched | C | 20259 | T | 15 | 0,1544 | 0,1548 | 0,0005 | 0,0002 | eq_cov | orf1ab | TTC | Phe | TTT | Phe | nsp15 |
| C29679T | enriched | C | 29679 | T | 13 | 0,1247 | 0,1250 | 0,0004 | 0,0003 | dep_pref | intergenic | - | - | - | - | intergenic |

**Table S11.**

Coordinates of mutated amino acids coupled with nucleotide mutations present in enriched *k*-mers identified in the bat dataset.

| mutation | type | ref_nucleotide | ref_pos | mut_nucleotide | n_kmers | max_bat_RMSE | agg_mutation_freq | early_agg_mutation_freq | group | gene | ref_codon | mut_codon | Protein |
|---|---|---|---|---|---|---|---|---|---|---|---|---|---|
| C230T | enriched | C | 230 | T | 15 | 0,4382 | 0,6388 | 0,0000 | dep_pref | intergenic | - | - | - |
| T232C | enriched | T | 232 | C | 15 | 0,4382 | 0,6654 | 0,0000 | dep_pref | intergenic | - | - | - |
| C241T | depleted | C | 241 | T | 15 | 0,1329 | 0,0152 | 0,1884 | dep_pref | intergenic | - | - | - |
| C280T | enriched | C | 280 | T | 7 | 0,4275 | 0,6046 | 0,0000 | dep_pref | orf1ab | GTC | GTT | nsp1 |
| A691T | enriched | A | 691 | T | 8 | 0,4356 | 0,6426 | 0,0000 | dep_pref | orf1ab | TCA | TCT | nsp1 |
| T693A | enriched | T | 693 | A | 9 | 0,4356 | 0,6502 | 0,0000 | dep_pref | orf1ab | TTT | TAT | nsp1 |
| C703T | enriched | C | 703 | T | 7 | 0,4164 | 0,6426 | 0,0005 | dep_pref | orf1ab | GGC | GGT | nsp1 |
| A761G | enriched | A | 761 | G | 9 | 0,3549 | 0,6920 | 0,0000 | dep_pref | orf1ab | AGC | GGC | nsp1 |
| A1124C | enriched | A | 1124 | C | 10 | 0,3845 | 0,5665 | 0,0000 | dep_pref | orf1ab | AGG | CGG | nsp2 |
| G1126T | enriched | G | 1126 | T | 12 | 0,3845 | 0,5703 | 0,0000 | dep_pref | orf1ab | AGG | AGT | nsp2 |
| A1540T | enriched | A | 1540 | T | 9 | 0,3683 | 0,5970 | 0,0000 | dep_pref | orf1ab | CCA | CCT | nsp2 |
| C3037T | depleted | C | 3037 | T | 15 | 0,1312 | 0,0342 | 0,1932 | dep_pref | orf1ab | TTC | TTT | nsp3 |
| C8782T | depleted | C | 8782 | T | 15 | 0,1281 | 0,0837 | 0,2307 | dep_pref | orf1ab | AGC | AGT | nsp4 |
| C10594A | enriched | C | 10594 | A | 9 | 0,4248 | 0,6046 | 0,0000 | dep_pref | orf1ab | AAC | AAA | 3CL-PRO |
| C10789T | enriched | C | 10789 | T | 9 | 0,3757 | 0,5475 | 0,0010 | dep_pref | orf1ab | GAC | GAT | 3CL-PRO |
| A10888T | enriched | A | 10888 | T | 9 | 0,4194 | 0,5932 | 0,0000 | dep_pref | orf1ab | GGA | GGT | 3CL-PRO |
| G11083T | depleted | G | 11083 | T | 15 | 0,1611 | 0,0000 | 0,2278 | dep_ex | orf1ab | TTG | TTT | nsp6 |
| G12205A | enriched | G | 12205 | A | 7 | 0,3872 | 0,6046 | 0,0000 | dep_pref | orf1ab | TTG | TTA | nsp8 |
| G12211A | enriched | G | 12211 | A | 13 | 0,4221 | 0,6540 | 0,0000 | dep_pref | orf1ab | AAG | AAA | nsp8 |
| A12235G | enriched | A | 12235 | G | 15 | 0,4624 | 0,6730 | 0,0000 | eq_cov | orf1ab | GAA | GAG | nsp8 |
| A12250T | enriched | A | 12250 | T | 12 | 0,3898 | 0,5856 | 0,0000 | dep_pref | orf1ab | GCA | GCT | nsp8 |

| | | | | | | | | | | | | | |
|---|---|---|---|---|---|---|---|---|---|---|---|---|---|
| T12304C | enriched | T | 12304 | C | 16 | 0,5001 | 0,7262 | 0,0000 | enr_pref | orf1ab | TAT | TAC | nsp8 |
| T12313A | enriched | T | 12313 | A | 9 | 0,3710 | 0,5285 | 0,0000 | dep_pref | orf1ab | GCT | GCA | nsp8 |
| T12340A | enriched | T | 12340 | A | 9 | 0,3925 | 0,5627 | 0,0000 | dep_pref | orf1ab | GTT | GTA | nsp8 |
| C12400T | enriched | C | 12400 | T | 15 | 0,4571 | 0,6616 | 0,0000 | eq_cov | orf1ab | CTC | CTT | nsp8 |
| A12634T | enriched | A | 12634 | T | 9 | 0,4651 | 0,6616 | 0,0000 | dep_pref | orf1ab | GCA | GCT | nsp8 |
| T12680C | enriched | T | 12680 | C | 13 | 0,4356 | 0,6388 | 0,0000 | dep_pref | orf1ab | TTA | CTA | nsp8 |
| G12694A | enriched | G | 12694 | A | 9 | 0,4087 | 0,5856 | 0,0000 | dep_pref | orf1ab | GAG | GAA | nsp9 |
| T12748A | enriched | T | 12748 | A | 9 | 0,4544 | 0,6426 | 0,0000 | dep_pref | orf1ab | ACT | ACA | nsp9 |
| C12754T | enriched | C | 12754 | T | 15 | 0,4544 | 0,6502 | 0,0000 | eq_cov | orf1ab | TGC | TGT | nsp9 |
| C13009T | enriched | C | 13009 | T | 15 | 0,4382 | 0,6274 | 0,0000 | eq_cov | orf1ab | GCC | GCT | nsp9 |
| A13021T | enriched | A | 13021 | T | 9 | 0,3952 | 0,5703 | 0,0000 | dep_pref | orf1ab | GTA | GTT | nsp9 |
| G13126A | enriched | G | 13126 | A | 10 | 0,4571 | 0,6768 | 0,0000 | dep_pref | orf1ab | GGG | GGA | nsp10 |
| A13327C | enriched | A | 13327 | C | 12 | 0,3979 | 0,5970 | 0,0000 | dep_pref | orf1ab | ACA | ACC | nsp10 |
| T13345A | enriched | T | 13345 | A | 7 | 0,3657 | 0,5627 | 0,0000 | dep_pref | orf1ab | CCT | CCA | nsp10 |
| T13387A | enriched | T | 13387 | A | 9 | 0,3845 | 0,5703 | 0,0000 | dep_pref | orf1ab | GGT | GGA | nsp10 |
| C13436A | enriched | C | 13436 | A | 8 | 0,4033 | 0,5703 | 0,0000 | dep_pref | orf1ab | CTT | ATT | nsp10 |
| T13521C | enriched | T | 13521 | C | 12 | 0,4191 | 0,5932 | 0,0005 | dep_pref | orf1ab | AGT | AGC | RDRP |
| C13548T | enriched | C | 13548 | T | 9 | 0,4490 | 0,6806 | 0,0000 | dep_pref | orf1ab | GAC | GAT | RDRP |
| T13578A | enriched | T | 13578 | A | 15 | 0,4786 | 0,6844 | 0,0000 | eq_cov | orf1ab | GCT | GCA | RDRP |
| A13581G | enriched | A | 13581 | G | 15 | 0,4786 | 0,6844 | 0,0000 | eq_cov | orf1ab | AAA | AAG | RDRP |
| T13599C | enriched | T | 13599 | C | 12 | 0,4140 | 0,6578 | 0,0000 | dep_pref | orf1ab | TGT | TGC | RDRP |
| T13602C | enriched | T | 13602 | C | 9 | 0,3764 | 0,6046 | 0,0000 | dep_pref | orf1ab | TGT | TGC | RDRP |
| C13743T | enriched | C | 13743 | T | 7 | 0,4356 | 0,6274 | 0,0000 | dep_pref | orf1ab | TTC | TTT | RDRP |
| T13746C | enriched | T | 13746 | C | 10 | 0,4356 | 0,6388 | 0,0000 | dep_pref | orf1ab | TTT | TTC | RDRP |
| A13756G | enriched | A | 13756 | G | 13 | 0,3898 | 0,5665 | 0,0000 | dep_pref | orf1ab | ATA | GTA | RDRP |

| | | | | | | | | | | | | | |
|---|---|---|---|---|---|---|---|---|---|---|---|---|---|
| C13761T | enriched | C | 13761 | T | 15 | 0,4140 | 0,5856 | 0,0000 | eq_cov | orf1ab | GAC | GAT | RDRP |
| A13917G | enriched | A | 13917 | G | 12 | 0,4329 | 0,6996 | 0,0000 | dep_pref | orf1ab | AAA | AAG | RDRP |
| C13923T | enriched | C | 13923 | T | 9 | 0,4329 | 0,6540 | 0,0000 | dep_pref | orf1ab | GAC | GAT | RDRP |
| G14137A | enriched | G | 14137 | A | 8 | 0,4194 | 0,6160 | 0,0000 | dep_pref | orf1ab | GTT | ATT | RDRP |
| G14335A | enriched | G | 14335 | A | 10 | 0,4087 | 0,5856 | 0,0000 | dep_pref | orf1ab | GTT | ATT | RDRP |
| C14408T | depleted | C | 14408 | T | 15 | 0,1281 | 0,0000 | 0,1812 | dep_ex | orf1ab | CCT | CTT | RDRP |
| T14496A | enriched | T | 14496 | A | 9 | 0,3898 | 0,6160 | 0,0000 | dep_pref | orf1ab | GGT | GGA | RDRP |
| G14622A | enriched | G | 14622 | A | 9 | 0,3898 | 0,5932 | 0,0000 | dep_pref | orf1ab | ACG | ACA | RDRP |
| C14697T | enriched | C | 14697 | T | 12 | 0,4651 | 0,6692 | 0,0000 | dep_pref | orf1ab | TTC | TTT | RDRP |
| T14749C | enriched | T | 14749 | C | 9 | 0,3657 | 0,5285 | 0,0000 | dep_pref | orf1ab | TTA | CTA | RDRP |
| T14778C | enriched | T | 14778 | C | 15 | 0,4705 | 0,7034 | 0,0000 | eq_cov | orf1ab | GGT | GGC | RDRP |
| C14793T | enriched | C | 14793 | T | 15 | 0,3818 | 0,5551 | 0,0000 | eq_cov | orf1ab | AGC | AGT | RDRP |
| C14805T | enriched | C | 14805 | T | 12 | 0,3818 | 0,6160 | 0,0778 | dep_pref | orf1ab | TAC | TAT | RDRP |
| T14808C | enriched | T | 14808 | C | 9 | 0,1216 | 0,1216 | 0,0000 | Unknown | orf1ab | TAT | TAC | nsp12 |
| A14820G | enriched | A | 14820 | G | 15 | 0,4275 | 0,6502 | 0,0000 | eq_cov | orf1ab | CTA | CTG | RDRP |
| A14847C | enriched | A | 14847 | C | 11 | 0,4140 | 0,6464 | 0,0000 | dep_pref | orf1ab | CTA | CTC | RDRP |
| T14853C | enriched | T | 14853 | C | 14 | 0,4140 | 0,6008 | 0,0000 | dep_pref | orf1ab | TTT | TTC | RDRP |
| A15159G | enriched | A | 15159 | G | 9 | 0,4356 | 0,6160 | 0,0000 | dep_pref | orf1ab | CAA | CAG | RDRP |
| T15258C | enriched | T | 15258 | C | 15 | 0,4786 | 0,7452 | 0,0000 | eq_cov | orf1ab | TAT | TAC | RDRP |
| T15309C | enriched | T | 15309 | C | 12 | 0,4033 | 0,5703 | 0,0000 | dep_pref | orf1ab | GAT | GAC | RDRP |
| A15435G | enriched | A | 15435 | G | 9 | 0,4517 | 0,6388 | 0,0000 | dep_pref | orf1ab | GAA | GAG | RDRP |
| C15763T | enriched | C | 15763 | T | 9 | 0,5779 | 0,6882 | 0,0000 | Unknown | orf1ab | CTA | TTA | nsp12 |
| G15768A | enriched | G | 15768 | A | 11 | 0,4087 | 0,6198 | 0,0000 | dep_pref | orf1ab | GTG | GTA | RDRP |
| A15777T | enriched | A | 15777 | T | 8 | 0,4060 | 0,5932 | 0,0000 | dep_pref | orf1ab | ATA | ATT | RDRP |
| G15906A | enriched | G | 15906 | A | 9 | 0,4813 | 0,6844 | 0,0000 | dep_pref | orf1ab | CAG | CAA | RDRP |

| | | | | | | | | | | | | | |
|---|---|---|---|---|---|---|---|---|---|---|---|---|---|
| G16221A | enriched | G | 16221 | A | 9 | 0,4248 | 0,6274 | 0,0000 | dep_pref | orf1ab | CCG | CCA | RDRP |
| G16377C | enriched | G | 16377 | C | 15 | 0,4490 | 0,7034 | 0,0000 | eq_cov | orf1ab | CCG | CCC | Hel |
| A16497T | enriched | A | 16497 | T | 9 | 0,4544 | 0,6958 | 0,0000 | enr_ex | orf1ab | GGA | GGT | Hel |
| T16590C | enriched | T | 16590 | C | 7 | 0,4087 | 0,5856 | 0,0000 | dep_pref | orf1ab | GGT | GGC | Hel |
| T16692C | enriched | T | 16692 | C | 12 | 0,4544 | 0,7034 | 0,0000 | dep_pref | orf1ab | GCT | GCC | Hel |
| C16806T | enriched | C | 16806 | T | 15 | 0,4221 | 0,6350 | 0,0000 | eq_cov | orf1ab | AAC | AAT | Hel |
| A16818G | enriched | A | 16818 | G | 12 | 0,3979 | 0,5970 | 0,0000 | dep_pref | orf1ab | CAA | CAG | Hel |
| A16821T | enriched | A | 16821 | T | 9 | 0,3979 | 0,5627 | 0,0000 | dep_pref | orf1ab | ATA | ATT | Hel |
| T16866G | enriched | T | 16866 | G | 12 | 0,4248 | 0,6388 | 0,0000 | dep_pref | orf1ab | GTT | GTG | Hel |
| C16870A | enriched | C | 16870 | A | 8 | 0,4248 | 0,6008 | 0,0000 | dep_pref | orf1ab | CGA | AGA | Hel |
| A16893G | enriched | A | 16893 | G | 12 | 0,3818 | 0,6008 | 0,0000 | dep_pref | orf1ab | TTA | TTG | Hel |
| T16908C | enriched | T | 16908 | C | 8 | 0,6008 | 0,7072 | 0,0000 | Unknown | orf1ab | TAT | TAC | nsp13 |
| T17514C | enriched | T | 17514 | C | 8 | 0,3791 | 0,5551 | 0,0000 | dep_pref | orf1ab | TGT | TGC | Hel |
| T17529A | enriched | T | 17529 | A | 9 | 0,4409 | 0,6236 | 0,0000 | dep_pref | orf1ab | ACT | ACA | Hel |
| C18388T | enriched | C | 18388 | T | 7 | 0,3603 | 0,5171 | 0,0000 | dep_pref | orf1ab | CTA | TTA | ExoN |
| C18486T | enriched | C | 18486 | T | 12 | 0,4382 | 0,7490 | 0,0000 | dep_pref | orf1ab | CTC | CTT | ExoN |
| A18861T | enriched | A | 18861 | T | 15 | 0,4302 | 0,6388 | 0,0000 | eq_cov | orf1ab | GCA | GCT | ExoN |
| C18888T | enriched | C | 18888 | T | 9 | 0,3979 | 0,6084 | 0,0000 | dep_pref | orf1ab | CAC | CAT | ExoN |
| A19767T | enriched | A | 19767 | T | 8 | 0,3952 | 0,5627 | 0,0000 | dep_pref | orf1ab | TTA | TTT | nsp15 |
| G20131T | enriched | G | 20131 | T | 12 | 0,4114 | 0,6350 | 0,0000 | dep_pref | orf1ab | GCC | TCC | nsp15 |
| C20133A | enriched | C | 20133 | A | 12 | 0,4194 | 0,6046 | 0,0000 | dep_pref | orf1ab | GCC | GCA | nsp15 |
| A20742G | enriched | A | 20742 | G | 11 | 0,4140 | 0,6236 | 0,0000 | dep_pref | orf1ab | CAA | CAG | nsp16 |
| C20775A | enriched | C | 20775 | A | 11 | 0,4167 | 0,6008 | 0,0000 | dep_pref | orf1ab | GGC | GGA | nsp16 |
| A23403G | depleted | A | 23403 | G | 15 | 0,1356 | 0,0000 | 0,1917 | dep_pref | spike | GAT | GGT | S |
| T26130A | enriched | T | 26130 | A | 11 | 0,4624 | 0,6540 | 0,0000 | dep_pref | orf3a | ATT | ATA | ORF3 |

| | | | | | | | | | | | | | |
|---|---|---|---|---|---|---|---|---|---|---|---|---|---|
| G26144T | depleted | G | 26144 | T | 15 | 0,1227 | 0,0000 | 0,1735 | dep_pref | orf3a | GGT | GTT | ORF3 |
| T26160A | enriched | T | 26160 | A | 12 | 0,4033 | 0,6312 | 0,0000 | dep_pref | orf3a | GTT | GTA | ORF3 |
| T26168C | enriched | T | 26168 | C | 7 | 0,3925 | 0,5703 | 0,0000 | dep_pref | orf3a | GTA | GCA | ORF3 |
| A26190G | enriched | A | 26190 | G | 15 | 0,4490 | 0,6730 | 0,0000 | eq_cov | orf3a | GAA | GAG | ORF3 |
| T26331C | enriched | T | 26331 | C | 15 | 0,3925 | 0,5894 | 0,0000 | eq_cov | e | GTT | GTC | E |
| C26395T | enriched | C | 26395 | T | 11 | 0,4490 | 0,6654 | 0,0000 | dep_pref | e | CTT | TTT | E |
| T26397A | enriched | T | 26397 | A | 9 | 0,4409 | 0,6540 | 0,0000 | dep_pref | e | CTT | CTA | E |
| A26482C | enriched | A | 26482 | C | 8 | 0,4813 | 0,6806 | 0,0000 | dep_pref | intergenic | - | - | - |
| A26693G | enriched | A | 26693 | G | 7 | 0,4087 | 0,5856 | 0,0000 | dep_pref | m | TTA | TTG | M |
| T26714C | enriched | T | 26714 | C | 8 | 0,4651 | 0,6730 | 0,0000 | dep_pref | m | TGT | TGC | M |
| A26741T | enriched | A | 26741 | T | 9 | 0,4140 | 0,6046 | 0,0000 | dep_pref | m | ATA | ATT | M |
| C26801T | enriched | C | 26801 | T | 10 | 0,4755 | 0,6730 | 0,0005 | dep_pref | m | CTC | CTT | M |
| G27065A | enriched | G | 27065 | A | 11 | 0,4194 | 0,6046 | 0,0000 | dep_pref | m | TTG | TTA | M |
| T27071G | enriched | T | 27071 | G | 10 | 0,3603 | 0,5247 | 0,0000 | dep_pref | m | GCT | GCG | M |
| G27112A | enriched | G | 27112 | A | 8 | 0,4275 | 0,6046 | 0,0000 | dep_pref | m | AGT | AAT | M |
| T27113C | enriched | T | 27113 | C | 7 | 0,4167 | 0,5894 | 0,0000 | dep_pref | m | AGT | AGC | M |
| G27163A | enriched | G | 27163 | A | 7 | 0,4678 | 0,7072 | 0,0000 | dep_pref | m | AGT | AAT | M |
| T27164C | enriched | T | 27164 | C | 8 | 0,4705 | 0,7110 | 0,0000 | dep_pref | m | AGT | AGC | M |
| T27182A | enriched | T | 27182 | A | 15 | 0,4893 | 0,7034 | 0,0000 | eq_cov | m | CTT | CTA | M |
| C27725T | enriched | C | 27725 | T | 12 | 0,4060 | 0,6198 | 0,0000 | dep_pref | orf7a | ACA | ATA | ORF7a |
| T28144C | depleted | T | 28144 | C | 15 | 0,1497 | 0,0342 | 0,2345 | dep_ex | orf8 | TTA | TCA | ORF8 |
| G28321A | enriched | G | 28321 | A | 10 | 0,4624 | 0,6540 | 0,0000 | dep_pref | n | ACG | ACA | N |
| C28435A | enriched | C | 28435 | A | 15 | 0,4194 | 0,5932 | 0,0000 | eq_cov | n | ACC | ACA | N |
| A28447G | enriched | A | 28447 | G | 10 | 0,4140 | 0,5894 | 0,0000 | dep_pref | n | CAA | CAG | N |
| A28547C | enriched | A | 28547 | C | 12 | 0,4544 | 0,6464 | 0,0000 | dep_pref | n | AGA | CGA | N |

| | | | | | | | | | | | | | |
|---|---|---|---|---|---|---|---|---|---|---|---|---|---|
| A28553G | enriched | A | 28553 | G | 15 | 0,4544 | 0,6464 | 0,0000 | eq_cov | n | ATT | GTT | N |
| T28570C | enriched | T | 28570 | C | 15 | 0,3683 | 0,5247 | 0,0000 | eq_cov | n | GGT | GGC | N |
| T28582G | enriched | T | 28582 | G | 9 | 0,3791 | 0,5703 | 0,0000 | dep_pref | n | GAT | GCG | N |
| G28621C | enriched | G | 28621 | C | 11 | 0,4033 | 0,5779 | 0,0000 | dep_pref | n | GGG | GGC | N |
| G28631T | enriched | G | 28631 | T | 7 | 0,3818 | 0,5627 | 0,0000 | dep_pref | n | GGA | TGA | N |
| C28657A | enriched | C | 28657 | A | 7 | 0,3683 | 0,5856 | 0,0000 | dep_pref | n | GAC | GAA | N |
| A28870T | enriched | A | 28870 | T | 11 | 0,4167 | 0,6160 | 0,0000 | dep_pref | n | CCA | CCT | N |
| C28887A | enriched | C | 28887 | A | 13 | 0,4060 | 0,6046 | 0,0000 | dep_pref | n | ACT | AAT | N |
| A28973G | enriched | A | 28973 | G | 13 | 0,4087 | 0,6502 | 0,0000 | dep_pref | n | ATG | GTG | N |
| G28975T | enriched | G | 28975 | T | 15 | 0,4087 | 0,6502 | 0,0019 | eq_cov | n | ATG | ATT | N |
| G29212A | enriched | G | 29212 | A | 12 | 0,6008 | 0,7452 | 0,0000 | Unknown | N | GCG | GCA | N |
| G29227A | enriched | G | 29227 | A | 15 | 0,4356 | 0,6198 | 0,0000 | eq_cov | n | TCG | TCA | N |
| A29306C | enriched | A | 29306 | C | 8 | 0,3818 | 0,5741 | 0,0000 | dep_pref | n | AAT | CAT | N |
| G29395A | enriched | G | 29395 | A | 11 | 0,4490 | 0,6502 | 0,0000 | enr_pref | n | AAG | AAA | N |
| G29399A | enriched | G | 29399 | A | 10 | 0,4114 | 0,5817 | 0,0000 | enr_ex | n | GCT | ACT | N |
| A29487G | enriched | A | 29487 | G | 7 | 0,3818 | 0,5399 | 0,0000 | enr_ex | n | AAA | AGA | N |
| C29518T | enriched | C | 29518 | T | 8 | 0,4487 | 0,6464 | 0,0005 | dep_pref | n | GAC | GAT | N |
| C29530A | enriched | C | 29530 | A | 8 | 0,4248 | 0,6008 | 0,0000 | dep_pref | n | GCC | GCA | N |
| C29541A | enriched | C | 29541 | A | 8 | 0,4060 | 0,5741 | 0,0000 | dep_pref | intergenic | - | - | - |
| A29542T | enriched | A | 29542 | T | 9 | 0,4060 | 0,5856 | 0,0000 | dep_pref | intergenic | - | - | - |
| A29567G | enriched | A | 29567 | G | 14 | 0,4490 | 0,6350 | 0,0000 | dep_pref | orf10 | ATA | GTA | ORF10 |
| T29582A | enriched | T | 29582 | A | 7 | 0,4114 | 0,6540 | 0,0000 | dep_pref | orf10 | TTT | ATT | ORF10 |
| T29597C | enriched | T | 29597 | C | 15 | 0,4463 | 0,6350 | 0,0000 | eq_cov | orf10 | TAT | CAT | ORF10 |
| A29649G | enriched | A | 29649 | G | 13 | 0,4356 | 0,6236 | 0,0000 | dep_pref | orf10 | GAT | GGT | ORF10 |
| G29651T | enriched | G | 29651 | T | 14 | 0,4436 | 0,6274 | 0,0000 | dep_pref | orf10 | GTA | TTA | ORF10 |

| | | | | | | | | | | | | | |
|---|---|---|---|---|---|---|---|---|---|---|---|---|---|
| G29688A | enriched | G | 29688 | A | 15 | 0,4167 | 0,6008 | 0,0000 | eq_cov | intergenic | - | - | - |
| T29758G | enriched | T | 29758 | G | 8 | 0,4705 | 0,6654 | 0,0000 | dep_pref | intergenic | - | - | - |

**Table S12.**

Recorded running times and peak memory usage for the datasets used in this study.
The computation times and memory usage were tested on a desktop PC with Intel Core i7-13700K 16 core 3.4 GHz CPU, 128 GB RAM and NVIDIA GeForce RTX 4090 GPU and a SSD hard drive.

| Dataset | Step | Computing time | Total memory required |
|---|---|---|---|
| Deer | k-mer extraction | 28 min 19 s | 313 MB |
| | Count matrix generation & rarity filter | 51 min 17 s | 23.5 GB |
| | Over-represented k-mers identification | 13 min 27 s | |
| | Re-examination of rare k-mers | 1 h 05 min 23 s | |
| | k-mer mapping & mutation identification | 21 min 55 s | |
| | Selection of linearly independent k-mers | 1 min 27 s | 1.5 GB |
| | k-mer importance for classification computation | 51 s | 2.6 GB |
| | Model architecture selection | 33 s | |
| | Model evaluation with bootstrapping | 9 min 20 s | |
| Bat | k-mer extraction | 2 min 03 s | 97 MB |
| | Count matrix generation & rarity filter | 6 min 50 s | 6.7 GB |
| | Over-represented k-mers identification | 50 s | |
| | Re-examination of rare k-mers | 47 min 24 s | |
| | k-mer mapping & mutation identification | 3 min 33 s | |
| | Selection of linearly independent k-mers | 2 h 33min 26 s | 84 GB |
| | k-mer importance for classification computation | 15 s | 12.2 GB |
| | Model architecture selection | 10 s | |
| | Model evaluation with bootstrapping | 6 min 20 s | |

**Table S13.**

Details of dense neural network (DNN) architecture, hyperparameters, and training.

| **Network architecture** | |
|---|---|
| **Hyperparameter** | **Value** |
| Number of hidden layers* | 2 |
| Number of nodes for hidden layers | [1024, 256] |
| Number of nodes for output layer | 1 |
| Activation function of hidden layers | GELU |
| Activation function of output layer | Sigmoid |
| Dropout ratio** | 0.3 |
| **Training** | |
| **Hyperparameter** | **Value** |
| Optimizer | AdamW |
| Learning rate | 1E-5 |
| Weight decay rate | 0.004 |
| Batch size | 8 |
| Loss function | Binary crossentropy |
| Class weights | balanced |
| Validation split | 0.1 |
| Training epochs | 50, with early stopping based on validation loss |

*Hidden layer weights were constrained to maximum norm of 3.
**Dropout was used before every hidden layer and before the output layer.
